# Identification of gene kinetics patterns improves RNA velocity and cell fate prediction using JuloVelo

**DOI:** 10.64898/2026.10.24.761531

**Authors:** Kuan-Chiun Tung, Yueh-Hua Tu, Kuan-Yi Hsieh, Hsueh-Fen Juan, Hsuan-Cheng Huang

## Abstract

Single-cell RNA velocity has transformed our capacity to predict future cell states from sequencing snapshots, but existing methods rely on constant or binary kinetic assumptions that fail to capture the heterogeneous splicing dynamics of cellular differentiation. Here we present JuloVelo, a deep learning framework that demonstrates that identifying gene kinetics patterns to initialize neural network training dramatically improves RNA velocity model performance. By pre-determining whether each gene follows a Circle, Complete induction, or Complete repression pattern across cellular differentiation, JuloVelo initializes training with pattern-matched, pre-trained network weights, effectively guiding the cell-specific kinetics optimization away from biologically implausible parameter regimes. Additionally, JuloVelo demonstrates that multiome datasets can also be processed and modeled by utilizing promoter chromatin accessibility to guide kinetics pre-determination, and integrates an *in silico* perturbation framework to simulate transcription factor modulation on cell fate. Validated across eight datasets spanning erythrogenesis, neurogenesis, and endocrinogenesis, JuloVelo consistently outperformed constant-rate and cell-specific models, resolving the entire five-lineage differentiation program in hippocampal neurogenesis. Multiome processing improved both intrinsic and extrinsic velocity stability, and JuloVelo reduced computational execution time by approximately 70% compared to cellDancer. JuloVelo is available as an open-source package at https://github.com/jhlabtw/JuloVelo.jl.

## 1 Introduction

Cell differentiation is governed by dynamic changes in gene expression that unfold continuously from progenitor to committed cell states. Single-cell RNA sequencing (scRNA-seq) has transformed our capacity to characterize these processes at cellular resolution [1–3], enabling the discovery of novel cell types, transitional intermediates, and lineage hierarchies across diverse biological systems. Trajectory inference algorithms such as Monocle [4], Palantir [5], and PAGA [6] reconstruct differentiation paths by modeling transcriptional similarity across cellular transitions. However, these approaches rely solely on snapshot expression profiles and cannot distinguish cells with similar transcriptional states but divergent future fates, limiting their ability to capture the directionality of developmental processes.

RNA velocity was introduced as a framework that exploits the kinetic relationship between unspliced pre-mRNA and mature spliced mRNA to predict the short-term future state of each cell [7]. Subsequent methods including scVelo [8] and UniTVelo [9] extended this approach through dynamical modeling of RNA metabolism, enabling robust reconstruction of differentiation directionality. However, all of these methods assume that transcription (α), splicing (β), and degradation (γ) rates are constant or binary across cells—an assumption that fails to capture the continuous, heterogeneous nature of gene regulation during differentiation [10]. Complex phenomena such as transcriptional boosting, multi-lineage branching, and cell cycle transitions involve kinetics that change substantially across developmental time, rendering constant-rate models fundamentally inadequate for such systems.

To overcome the constant kinetics limitation, recent deep learning approaches including DeepVelo [11] and cellDancer [12] introduced cell-specific kinetics modeling, allowing each cell to have distinct kinetic parameters inferred from local neighbor information. These methods provide greater flexibility for capturing regulatory heterogeneity. However, two key challenges remain. First, both methods initialize neural networks without explicit biological priors about each gene’s kinetic trajectory, which can lead to convergence toward kinetically inconsistent parameter estimates, particularly for genes exhibiting complex dynamic behaviors. Second, neither framework leverages the increasing availability of simultaneous chromatin accessibility measurements from multiome assays, which could provide independent constraints on transcriptional activity and improve inference accuracy.

Here we present JuloVelo, a neural network-based RNA velocity framework that addresses these limitations through gene kinetics trajectory identification. Building on cellDancer’s local, cell-specific inference paradigm [12], JuloVelo categorizes each gene into one of three kinetic patterns—Circle, Complete induction, or Complete repression—based on the relationship between unspliced and spliced RNA across the cellular state progression, and initializes training with pattern-matched pretrained networks. For multiome datasets, JuloVelo further demonstrates how promoter chromatin accessibility can be used as an additional constraint to guide kinetics predetermination. Additionally, JuloVelo provides an *in silico* perturbation framework that leverages inferred cell-specific kinetics to simulate transcription factor modulation and predict its effect on cell fate trajectories.

We validate JuloVelo across eight biological datasets spanning erythroid differentiation, hippocampal and pancreatic endocrinogenesis, skin and follicle development, and embryonic brain neurogenesis, as well as on simulated data with defined complex kinetics. Across these benchmarks, JuloVelo consistently outperformed scVelo, DeepVelo, and cellDancer, being the only method to correctly infer the complete differentiation trajectory in the mouse erythroid gastrulation dataset and to resolve all five differentiation branch lineages in hippocampus neurogenesis. Utilizing promoter chromatin accessibility during pre-determination improved both intrinsic and extrinsic velocity stability in multiome datasets, and JuloVelo reduced execution time by approximately 70% compared to cellDancer. Furthermore, cell-specific transcriptional rates estimated by JuloVelo provide superior identification of functionally relevant genes compared to conventional differential expression analysis, and *in silico* transcription factor perturbations reveal predictable and biologically interpretable shifts in cell fate trajectories.

In summary, JuloVelo represents a significant shift from static, curve-fitting descriptions of gene expression toward mechanistic, dynamic modeling of cellular decision-making. By establishing gene kinetics trajectory identification as a foundational biological prior for deep learning, JuloVelo provides a robust, scalable framework capable of resolving complex developmental branches and transcriptional boosting events that have historically confounded traditional velocity models. The capacity to guide neural network initialization with epigenomic data represents a major step toward unifying single-cell multi-omic profiling with biophysical kinetics. As comprehensive multi-omic atlases of development and disease continue to grow, JuloVelo’s combination of cell-specific kinetics inference, multi-omic dataset processing, and predictive *in silico* perturbations offers a powerful computational resource for dissecting and engineering cell fate trajectories in development, regeneration, and disease.

## 2 Results

### Overview of the JuloVelo framework

To address the limitations of constant-kinetics assumptions and unguided neural network initialization, we developed JuloVelo, a deep learning framework that introduces gene-level kinetics trajectory identification as a foundational biological prior for cell-specific RNA velocity inference (Fig. 1). The framework accepts single-cell unspliced and spliced RNA counts as primary inputs, with the optional incorporation of promoter chromatin accessibility profiles for multiome datasets (Fig. 1a). The workflow proceeds through three core stages. First, JuloVelo pre-determines gene-specific kinetics patterns across the differentiation progression, classifying each gene as Circle (exhibiting both induction and repression phases), Complete induction, or Complete repression (Fig. 1b). Second, these pre-determined patterns guide the assignment of pre-determined neural networks with pattern-matched weights, which are then fine-tuned to estimate cell-specific transcription (α), splicing (β), and degradation (γ) rates, preventing convergence to biologically implausible parameter regimes. Third, the estimated cell-specific parameters are used to project velocity vectors onto low-dimensional embeddings and reconstruct continuous developmental trajectories. For multiome datasets, promoter chromatin accessibility is integrated into the predetermination step as an additional prior to refine kinetics classification at loci with ambiguous RNA expression.

**Fig. 1.**
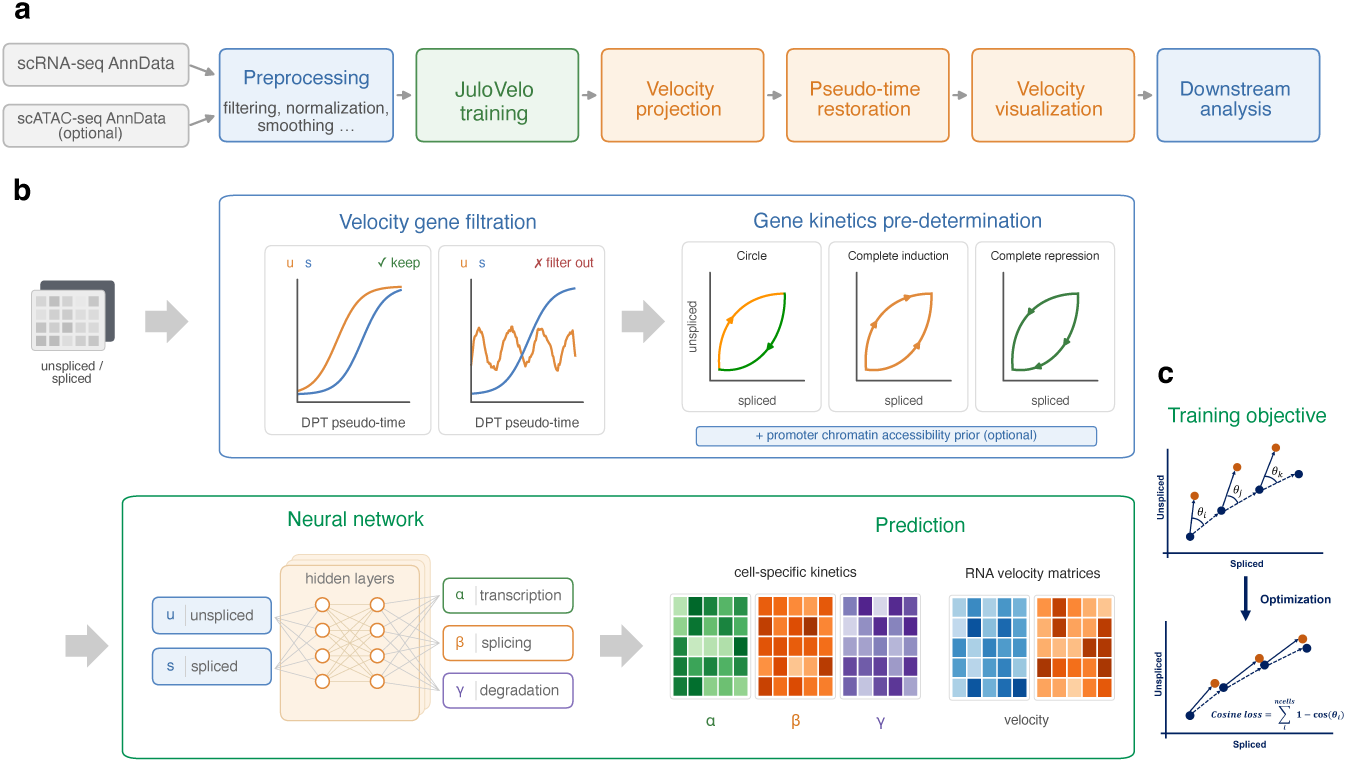
Overview of the JuloVelo framework. **a**. JuloVelo takes scRNA-seq AnnData containing unspliced and spliced RNA counts, with optional scATAC-seq AnnData for multiome datasets, and performs preprocessing, model training, velocity projection, pseudo-time restoration, velocity visualization, and downstream analysis. **b**. Prior to model training, genes are filtered according to their unspliced and spliced expression dynamics along diffusion pseudo-time (DPT), and gene kinetic trajectories are pre-determined as Circle, Complete induction, or Complete repression to initialize model training, optionally incorporating promoter chromatin accessibility. A neural network then uses unspliced and spliced RNA abundances to infer cell-specific transcription (*α*), splicing (*β*), and degradation (*γ*) rates and generate RNA velocity matrices. **c**. Model optimization aligns predicted RNA velocity directions with local cell-state transitions in the unspliced–spliced phase space using a cosine-based training objective.

### JuloVelo accurately infers complex gene kinetics in simulated data

We first validated JuloVelo’s capacity to accurately reconstruct velocity vector fields and estimate time-varying parameters on simulated datasets containing known ground-truth kinetics for four challenging scenarios: Circle, Transcriptional boosting, Multiple induction, and Multiple repression (Fig. 2a). As demonstrated by the projected velocity stream plots, JuloVelo correctly recovered the expected velocity vectors across all four kinetics regimes, capturing the outward flow from steady-state lines during induction and the inward flow during repression (Fig. 2b). Notably, for the Transcriptional boosting pattern—where the transcription rate sharply increases midprocess—JuloVelo successfully resolved both pre-boost and post-boost phases, with minor directionality deviations restricted to terminal cells due to sparse neighbor cell density. Furthermore, analysis of the estimated cell-specific parameters revealed that the inferred transcription rates (α) closely mirrored the true bimodal and boosting profiles, while degradation rates (γ) remained robustly below the theoretical maximum, consistent with post-transcriptional protective mechanisms (Fig. 2c–e). These synthetic benchmarks confirm that JuloVelo’s pattern-matched initialization reliably prevents the neural network from collapsing into non-functional parameter regimes, enabling accurate velocity recovery in complex biophysical systems.

**Fig. 2.**
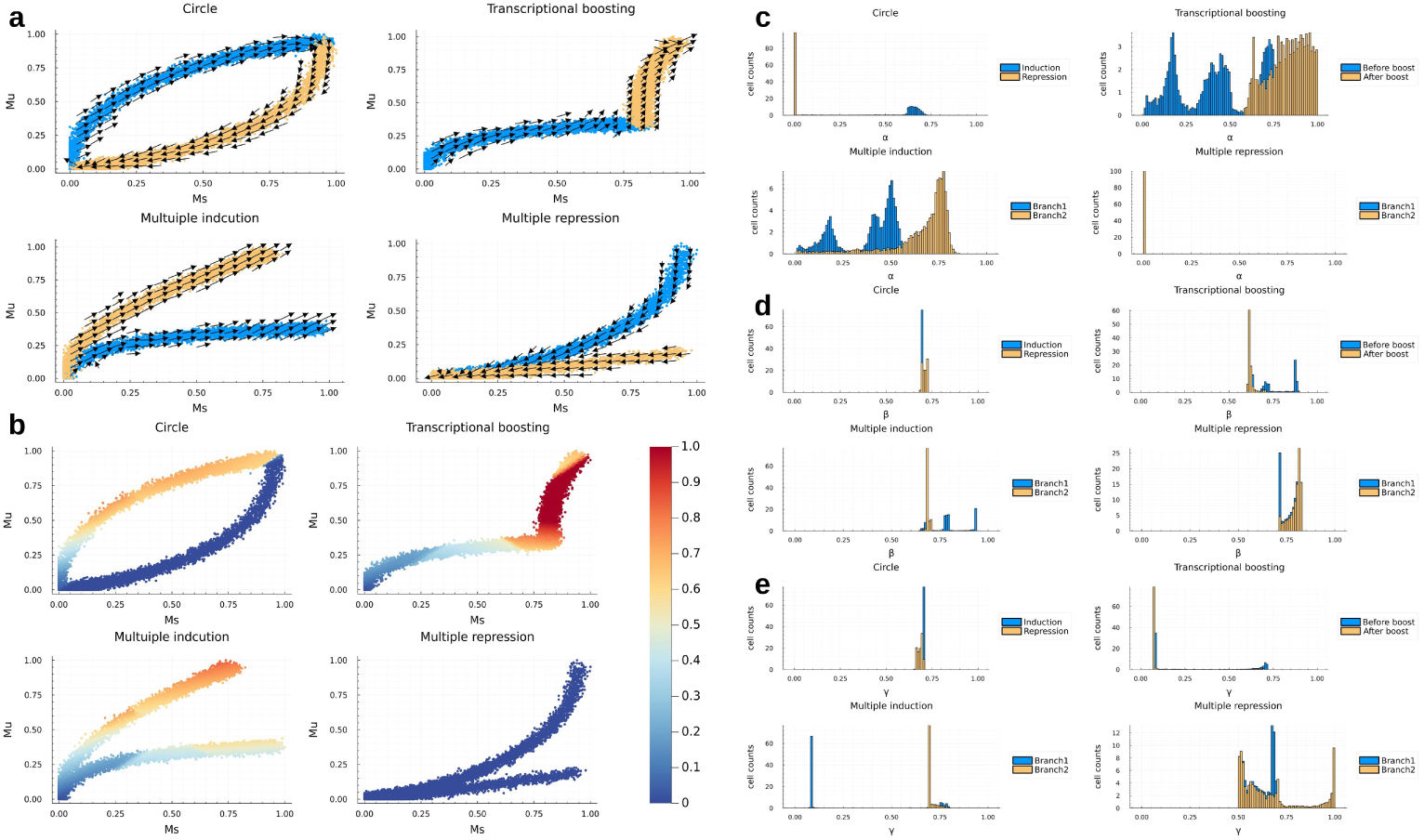
JuloVelo recovers complex gene kinetics in simulated data. Velocity flow and transcriptional-rate recovery across four challenging kinetics regimes—Circle, Transcriptional boosting, Multiple induction, and Multiple repression—demonstrating that pattern-matched initialization prevents collapse into non-functional parameter regimes (**a,b**), with inferred transcription rates (*α*) closely matching ground-truth profiles while degradation rates (*γ*) remain below the theoretical maximum (**c–e**).

### Gene kinetics trajectory identification enables accurate erythroid differentiation flow

To evaluate JuloVelo on a continuous developmental trajectory with known biological directionality, we benchmarked it against scVelo, DeepVelo, and cellDancer on a mouse erythroid gastrulation dataset featuring transcriptional boosting of key erythroid genes (Fig. 3). Visualized on the UMAP embedding, JuloVelo was the only method that correctly inferred a continuous, forward-directed differentiation flow from Blood progenitors 1 and 2, through transitional Erythroid 1 and 2 stages, to mature Erythroid 3 cells (Fig. 3a). In contrast, scVelo produced disjointed and backward flows, cellDancer failed to capture the progenitor transition, and DeepVelo exhibited a reversed vector field in the mature erythroid population. Consequently, JuloVelo correctly restored a continuous and smooth maturation gradient reflecting the true developmental progression (Fig. 3b). At the single-gene level, JuloVelo accurately reconstructed the cell-specific kinetics of key lineage drivers: *Hba-x* exhibited the expected transcriptional boosting surge in transcription rate (α) during mid-differentiation [13], whereas the early progenitor marker *Rspo3* showed a clear transition to a repression state upon cell-type commitment (Fig. 3c–f). This biological accuracy was further validated by a strong temporal concordance between the rolling mean of raw unspliced counts and the inferred cell-specific transcriptional rates across the differentiation progression (Fig. 3g). Finally, projection of the cell-specific kinetic parameters (α, β, γ) onto a principal component space revealed a continuous, low-dimensional trajectory that closely recapitulated the cell-type maturation path (Fig. 3h), demonstrating that JuloVelo’s cell-specific kinetics capture cellular differentiation as a continuous biophysical process [14].

**Fig. 3.**
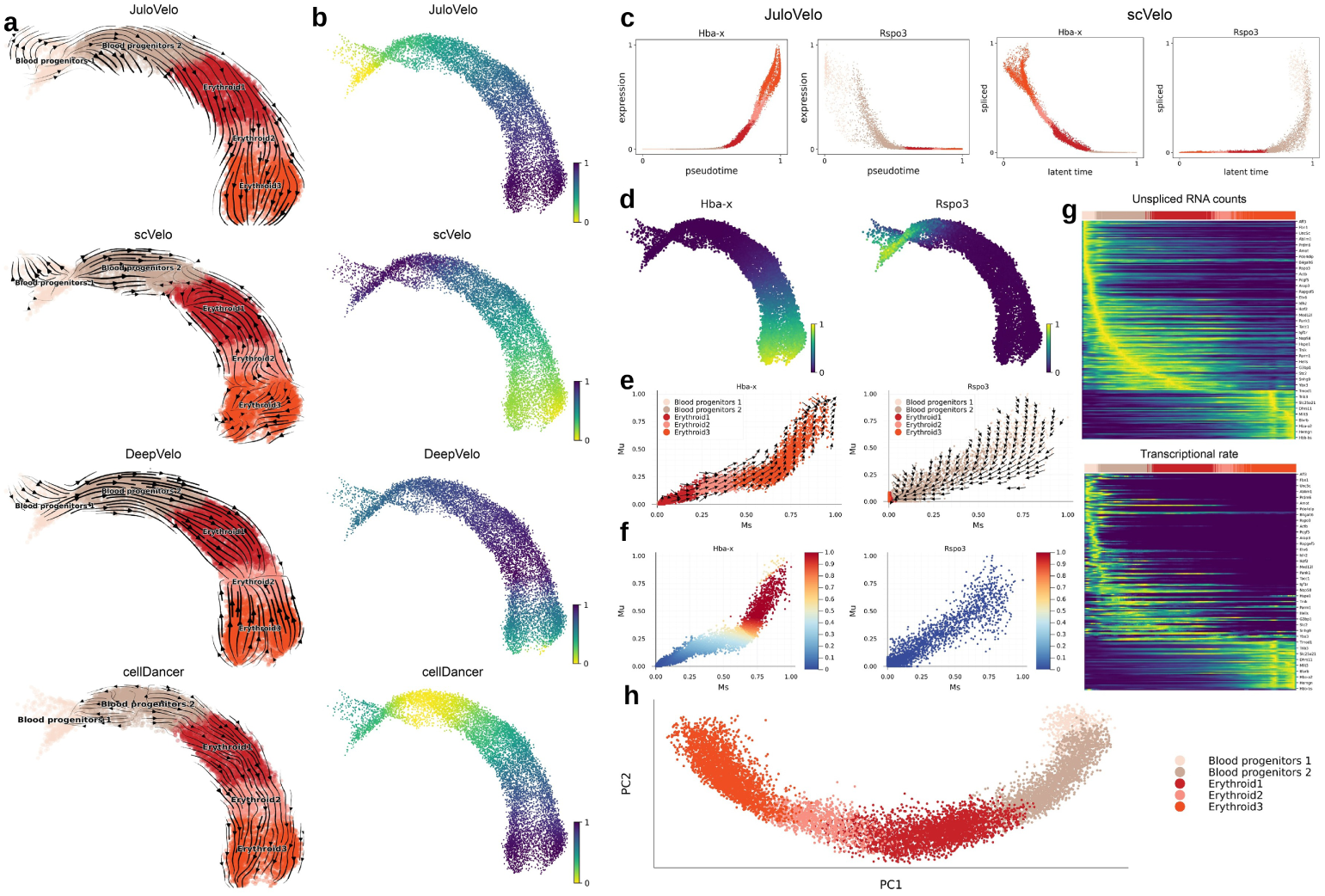
Gene kinetics trajectory identification enables accurate erythroid differentiation flow. On mouse erythroid gastrulation, JuloVelo is the only method that infers a continuous, forward-directed flow and a smooth maturation gradient, whereas scVelo, DeepVelo, and cellDancer produce disjointed, reversed, or absent transitions (**a,b**). At the single-gene level, JuloVelo reconstructs the transcriptional boosting of *Hba-x* and the repression of the early progenitor marker *Rspo3*, with cell-specific parameters projecting onto a continuous maturation path (**c–h**).

### JuloVelo resolves multi-lineage dynamics in hippocampus neurogenesis

We next tested JuloVelo’s capability to resolve complex branching trajectories using a mouse hippocampal neurogenesis dataset, which features five distinct differentiation lineages branching from neuron intermediate progenitor cells (nIPCs) (Fig. 4a). Among the evaluated models, JuloVelo was the only framework that successfully resolved all five developmental branches—including the oligodendrocyte, glial, and three distinct neuronal lineages—and recovered their correct developmental orderings (Fig. 4a,b). Specifically, scVelo produced completely reversed flows along the granule neuron lineage, cellDancer generated chaotic vectors in the transitional neuroblast stages, and DeepVelo failed at the critical CA1-Subiculum boundary. At the individual gene level, JuloVelo correctly categorized and modeled the distinct kinetics of lineage-specific markers, such as the circular induction-repression dynamics of the neuroblast marker *Dcx* [15], the exclusive induction of *Sphkap*, which encodes a sphingosine kinase 1-interacting A-kinase anchoring protein [16], along the granule neuron lineage, and the induction of *Pid1* along the oligodendrocyte lineage (Fig. 4i–k). Heatmaps of the unspliced RNA counts and corresponding inferred transcription rates (α) ordered along each of the five branches revealed highly concordant, branch-specific activation cascades, confirming that JuloVelo captures the underlying gene regulatory programs (Fig. 4c–g). This is further substantiated by the low-dimensional embedding of the cell-specific kinetics parameters, which faithfully recapitulated the multi-lineage topology of the dentate gyrus (Fig. 4h), proving that JuloVelo’s localized inference paradigm can robustly map complex, multi-branched developmental landscapes.

**Fig. 4.**
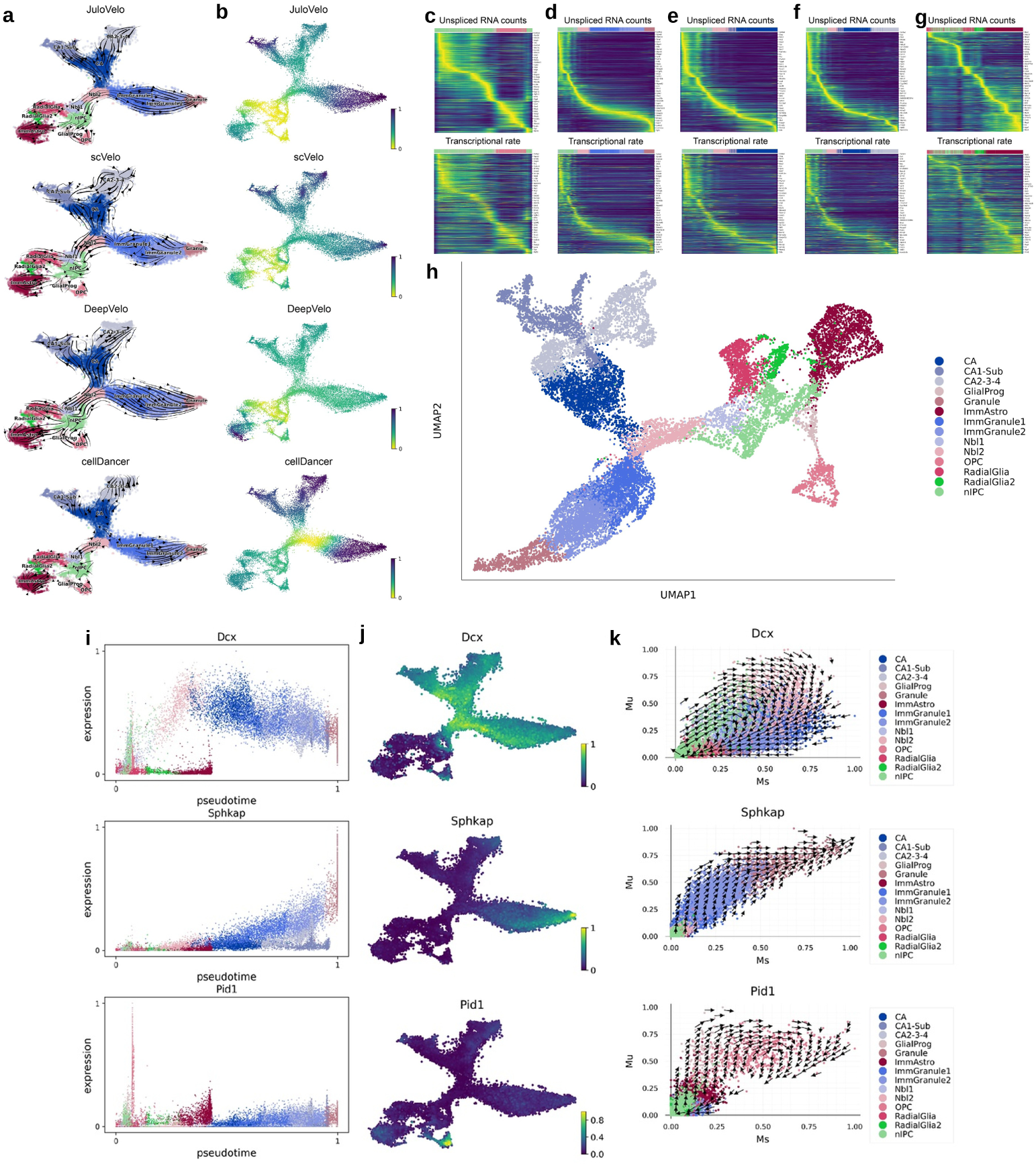
JuloVelo resolves multi-lineage dynamics in hippocampal neurogenesis. JuloVelo is the only framework that resolves all five differentiation branches and their correct orderings from neuron intermediate progenitor cells (**a,b**), with cell-specific kinetics reconstructing the multi-lineage topology of the dentate gyrus (**h**). Lineage-specific markers such as *Dcx*, *Sphkap*, and *Pid1* show concordant branch-specific activation cascades between unspliced RNA and inferred transcription rates (**c–g, i–k**).

### Cell-specific kinetics captures regulatory dynamics in pancreatic endocrinogenesis

We extended our evaluation to mouse pancreatic endocrinogenesis, a multi-lineage system where ductal progenitors differentiate into four distinct hormone-producing endocrine cell types (Fig. 5a). JuloVelo successfully mapped the continuous differentiation flow originating from Ductal cells, progressing through Ngn3-high endocrine progenitors and Pre-endocrine cells, and branching into mature alpha, beta, delta, and epsilon cell types (Fig. 5a,b). Interestingly, within the Ductal and Ngn3-low progenitor clusters JuloVelo captured a cyclic velocity flow corresponding to the S-to-G2M cell-cycle transition, consistent with the S-phase and G2M-phase scores estimated from cell-cycle gene expression (Fig. 5c,d). At the single-gene level, JuloVelo’s cell-specific kinetics successfully captured the transient transcriptional states that define these lineage decisions (Fig. 5e,f). For example, *Actn4* exhibited a clear repression state in pre-endocrine cells; while its steady-state mature spliced RNA remained highly detectable, JuloVelo correctly inferred that active transcription had already ceased. This demonstrates that JuloVelo can resolve temporal repression dynamics that are completely invisible to static expression-based analyses, highlighting the value of cell-specific kinetics for tracking active regulatory states.

**Fig. 5.**
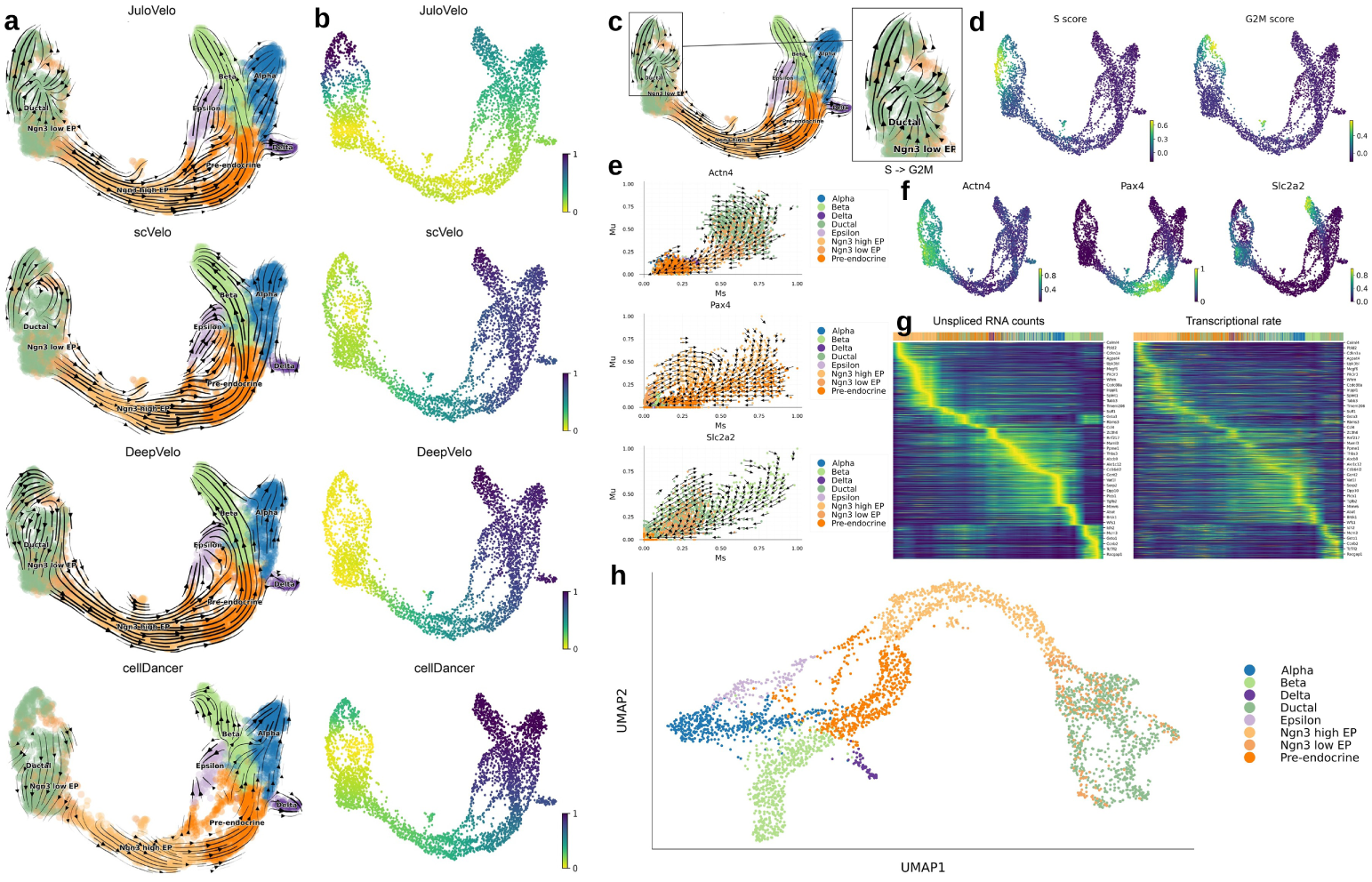
Cell-specific kinetics captures regulatory dynamics in pancreatic endocrinogenesis. **a**, Velocity streamlines inferred by JuloVelo, scVelo, DeepVelo, and cellDancer on the UMAP embedding, showing the continuous flow from Ductal progenitors through Ngn3-high endocrine pro-genitors (EP) and Pre-endocrine cells into mature alpha, beta, delta, and epsilon cells. **b**, Pseudo-time restored from the velocity inferred by each method. **c**, JuloVelo velocity flow in Ductal and Ngn3-low EP cells (inset), revealing a cyclic flow corresponding to the S-to-G2M cell-cycle transition. **d**, S-phase and G2M-phase scores estimated from the expression of cell-cycle genes. **e**, Unspliced–spliced phase portraits with JuloVelo velocity vectors for *Actn4*, *Pax4*, and *Slc2a2*, coloured by cell type. **f**, Spliced expression of *Actn4*, *Pax4*, and *Slc2a2* on the UMAP embedding; *Actn4* remains highly detectable in pre-endocrine cells although JuloVelo infers that its active transcription has ceased. **g**, Unspliced RNA counts (left) and JuloVelo-inferred transcription rates (right) of velocity genes ordered along pseudo-time show concordant patterns. **h**, UMAP of the cell-specific kinetic parameters (*α*, *β*, *γ*) inferred by JuloVelo, coloured by cell type.

### Performance on additional datasets

We further applied JuloVelo to two additional GEX datasets: human bone marrow hematopoiesis (5,292 cells across 9 cell types, after removing common lymphoid progenitors) and the mouse gastrulation subset to E7.5 (7,202 cells across 27 cell types). In human bone marrow, all four methods produced inconsistent flows within the HSC 1 and HSC 2 populations, and JuloVelo additionally showed unexpected flows in early HSC 1 and in immune cell types (Supplementary Fig. S1). Nevertheless, the HSC-expressed genes *CD44*, *TAOK3*, and *CELF2* displayed specific kinetics with opposite velocity directions in HSCs and immune cells, and the kinetics UMAP and the heatmaps of unspliced counts and transcription rates were concordant (Supplementary Fig. S2). In the gastrulation E7.5 dataset, the flows inferred by scVelo, DeepVelo, and cellDancer contained directional errors, whereas JuloVelo was the only method that recovered a reasonable pseudotime ordering (Supplementary Fig. S3). *Runx1* showed induction-only kinetics along erythroid differentiation, the kinetics UMAP reconstructed the full differentiation process, and unspliced counts and transcription rates were again concordant (Supplementary Fig. S4).

In the mouse skin and hair follicle dataset (6,436 cells across 5 cell types), in which rapidly proliferating transit-amplifying cells (TACs) differentiate into the inner root sheath (IRS) and the hair shaft cuticle, cortex, and medulla, JuloVelo inferred a reasonable flow and pseudotime starting from TAC-1 and terminating in the IRS, hair shaft, and medulla, although its flow within TAC-1 and TAC-2 was slightly irregular; the other three methods failed to recover a reasonable flow or pseudotime (Supplementary Fig. S5). TAC-1 and TAC-2 each contained several evenly represented Leiden clusters, suggesting unstable expression states that complicate flow inference for all methods. JuloVelo nonetheless captured the expected kinetics of the Medulla-expressed *Aldh1a3* (induction only), the TAC marker *Anln* (repression only), and *Adamts9* (early induction followed by repression), and inferred more non-linear expression dynamics along pseudotime than the ODE-based scVelo (Supplementary Fig. S6).

Finally, the mouse dentate gyrus neurogenesis dataset (2,930 cells across 14 cell types) revealed a limitation of local, neighbor-based velocity inference. On the full dataset, JuloVelo and cellDancer produced chaotic flows along the main granule and astrocyte lineages, whereas scVelo and DeepVelo inferred more continuous flows (Supplementary Fig. S7a). Phase portraits of *Tmeff2*, *Jph1*, and *Col19a1* showed that isolated populations such as oligodendrocytes and GABAergic interneurons distort the unspliced–spliced distribution of the main branch (Supplementary Fig. S8d). After these isolated populations were removed, JuloVelo recovered continuous flows and a continuous kinetics embedding of the main branch (Supplementary Figs. S7b and S8), and its intrinsic stability increased (Fig. 9e,f). As previously reported for scVelo [8], none of the four methods correctly inferred flow terminating in mature granule cells, whose transcriptional patterns closely resemble those of immature granule cells.

### Transcriptional rates outperform expression for identifying functional genes

Because JuloVelo estimates cell-specific transcription rates (α) directly, we hypothesized that comparing active transcriptional rates would yield more biologically relevant marker genes than comparing static expression levels, which suffer from mature mRNA retention. We performed differential transcriptional rate analysis (using JuloVelo’s cell-specific α) and conventional differential expression analysis across the pancreatic endocrine lineages (Fig. 6). Ranking genes by their cell-specific transcription rates (α) consistently identified a higher proportion of established, pathway-relevant endocrine regulators in beta, alpha, and delta cells compared to rankings based on static expression (Fig. 6a). For instance, genes actively driving beta-cell identity were ranked significantly higher by transcriptional rate, whereas genes that were biologically repressed but still retained mature transcripts were correctly filtered out (Fig. 6b). Consequently, downstream Gene Ontology (GO) [17] and KEGG pathway [18] enrichment analyses performed on differential transcriptional rate genes (DTRGs) yielded significantly more specific and statistically significant developmental pathways (Fig. 6c,d). These findings demonstrate that JuloVelo’s cell-specific parameters provide a superior quantitative foundation for identifying functional driver genes than standard expression profiling.

**Fig. 6.**
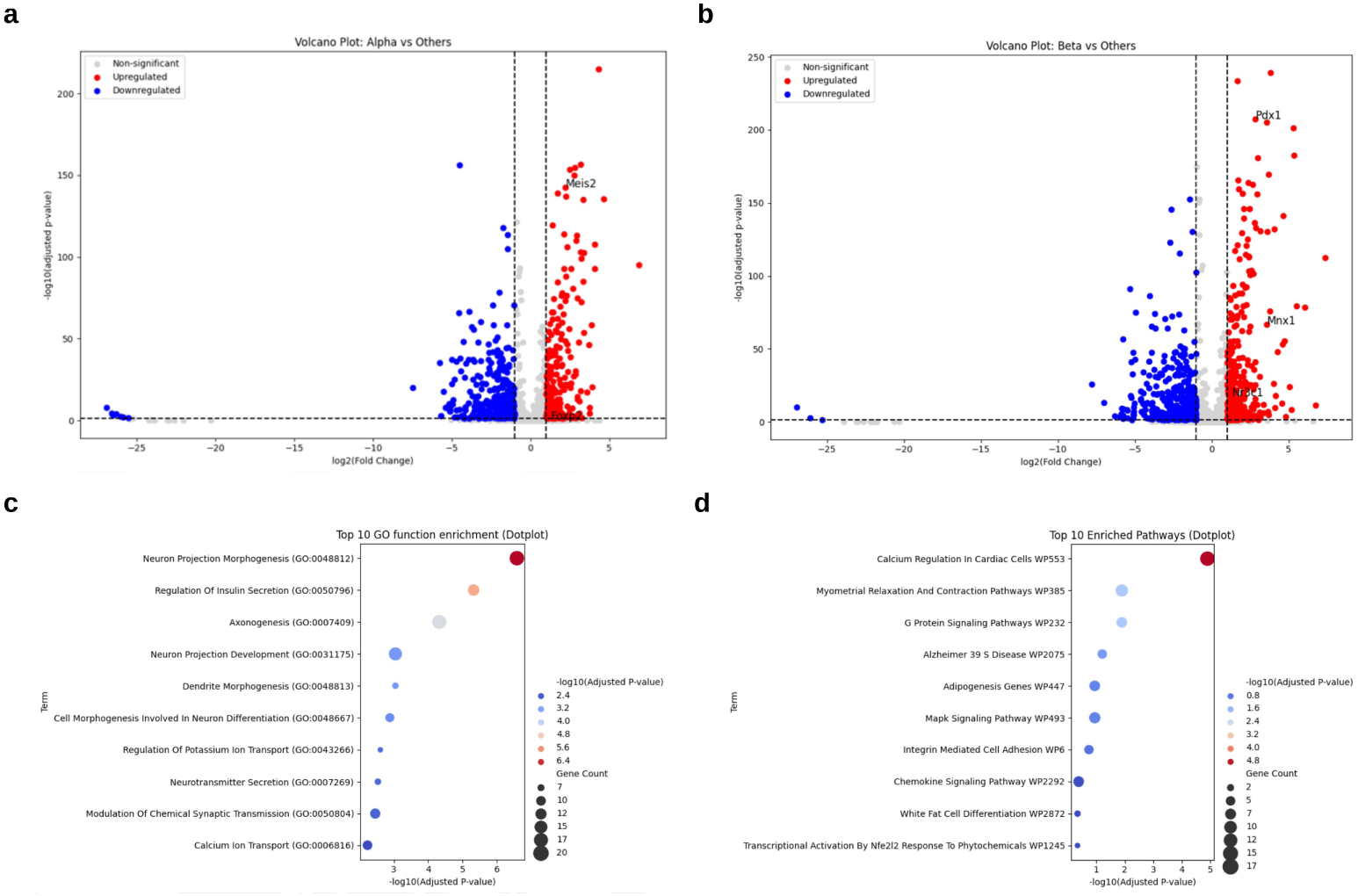
Transcriptional rates outperform expression for identifying functional genes. Conventional differential-expression analysis retains genes that are biologically repressed but still detectable as mature transcripts (**a,b**). Differential transcriptional rate analysis using JuloVelo’s cell-specific *α* (DTRGs) instead captures active regulatory programs, yielding Gene Ontology (**c**) and KEGG pathway (**d**) enrichments of higher specificity and statistical significance for lineage-specific transcription-factor targets.

### Processing multiome data improves velocity inference in embryonic brain

To determine whether utilizing epigenomic data during pre-determination resolves kinetically ambiguous loci, we evaluated JuloVelo on a Mouse Embryonic E18 Brain multiome dataset featuring parallel astrocyte and migrating neuron lineages (Fig. 7a). When run in RNA-only mode, all four tested methods exhibited substantial errors: scVelo failed to connect the V-SVZ and subplate, DeepVelo showed disjointed transitions, and cellDancer reversed the direction of mature deeper layer neurons (Fig. 7a,b). However, when we applied JuloVelo in multiome mode—utilizing promoter chromatin accessibility in the pre-determination step to guide kinetics patterns—it successfully restored a continuous, biologically correct differentiation flow connecting the upper and deeper cortical layers (Fig. 7c). This epigenomic guidance also led to a more biologically plausible root cell assignment and smoother differentiation gradients (Fig. 7d). At the single-gene level, JuloVelo multiome mode resolved key markers such as *Eomes* and *Gria2*, where promoter accessibility showed strong concordance with the onset of unspliced RNA induction (Fig. 7e–h). Crucially, for the deeper-layer marker *Tle4*, velocity vectors under RNA-only mode were highly unstable and circular, but became strongly directed and stable when guided by promoter chromatin accessibility (Fig. 7i,j). Quantitative evaluation confirmed that multiome processing significantly increased both intrinsic (within cell-type) and extrinsic (across boundaries) velocity stability compared to the RNA-only model (Fig. 9g), demonstrating that utilizing promoter chromatin accessibility to pre-determine kinetics patterns is highly effective for resolving transcriptomic ambiguities.

**Fig. 7.**
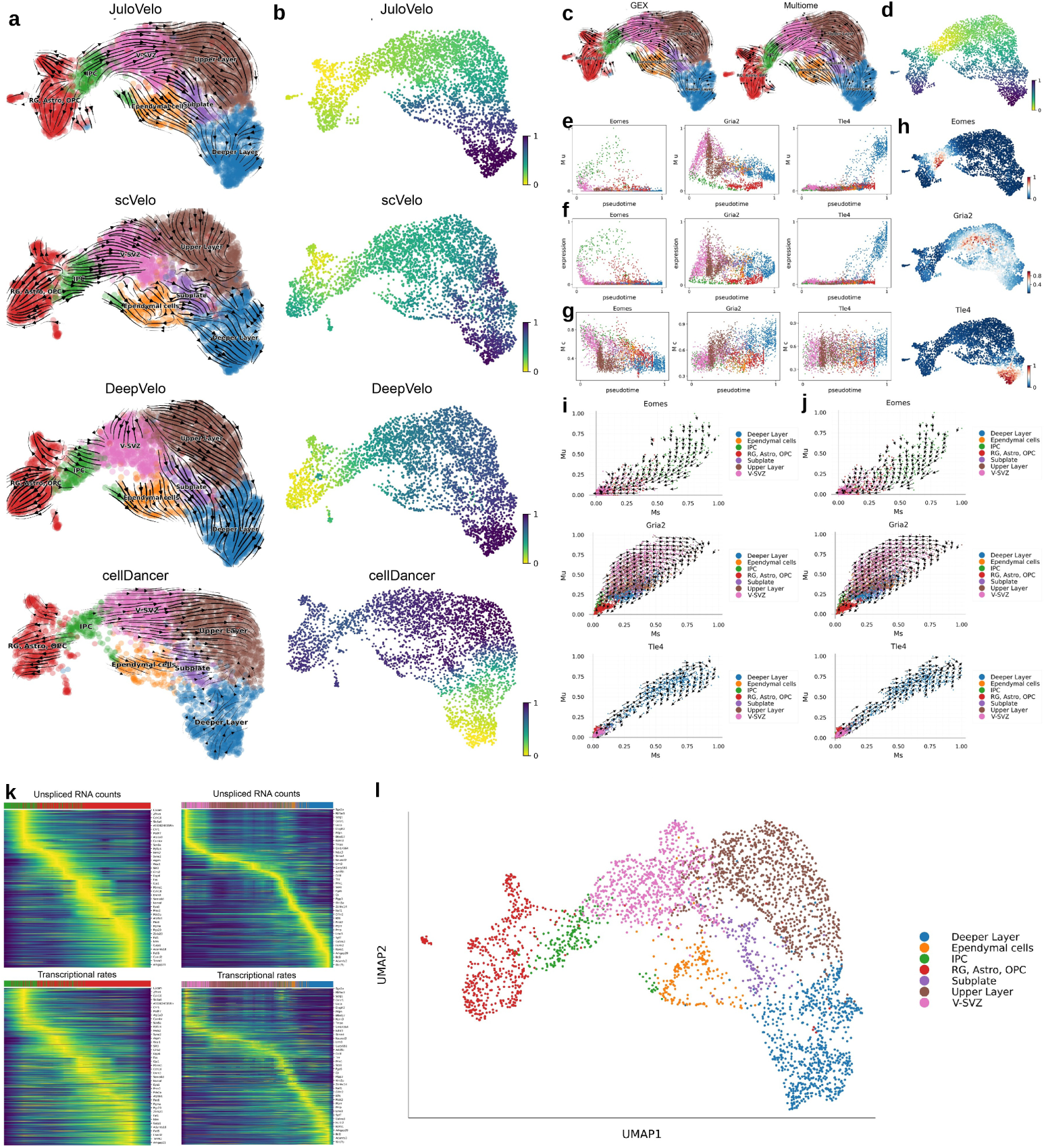
Processing multiome data improves velocity inference in embryonic brain. In a Mouse Embryonic E18 Brain multiome dataset, RNA-only inference produces disconnected or reversed flows across all tested methods (**a,b**). Incorporating promoter chromatin accessibility during kinetics pre-determination restores a continuous, biologically correct flow connecting the upper and deeper cortical layers (**c,d**), with single-gene markers such as *Eomes*, *Gria2*, and *Tle4* showing strong concordance between promoter accessibility and unspliced-RNA induction (**e–j**).

### In silico perturbation reveals TF-dependent cell fate trajectories

To demonstrate that JuloVelo’s cell-specific kinetics can be used to predict the down-stream regulatory consequences of transcription factor (TF) activity, we developed an *in silico* perturbation framework and applied it to pancreatic endocrinogenesis and hippocampal neurogenesis (Fig. 8). We first identified lineage-specific TFs and their downstream target regulons using pySCENIC activity scoring (AUCell). In pancreatic endocrinogenesis, we targeted the key beta-cell regulators Pdx1, Mnx1, and Nr3c1, and the alpha-cell drivers Meis2 and Foxp2. Without perturbation, most simulated pre-endocrine cells reached the beta-cell cluster (Fig. 8d). Knockdown simulations—where transcription rates (α) of target regulons were set to zero—of the beta-cell TFs in pre-endocrine progenitor cells resulted in simulated trajectories that deviated sharply from the expected beta-cell lineage, instead converging onto alternative alpha or delta-cell clusters (Fig. 8a). Conversely, overexpression simulations of these same TFs reinforced commitment toward the beta-cell fate. Similar lineage disruptions were observed upon knockdown of Meis2 and Foxp2 in alpha cells (Fig. 8b). To verify generalizability, we simulated knockdown of Zmiz1 in hippocampal Imm-Granule1 cells and Klf7 in CA cells, which successfully recapitulated developmental blockades and alternative lineage branching (Fig. 8c). These results establish that the cell-specific kinetics inferred by JuloVelo are mechanistically informative, enabling predictive, time-resolved simulations of cellular engineering and fate perturbation.

**Fig. 8.**
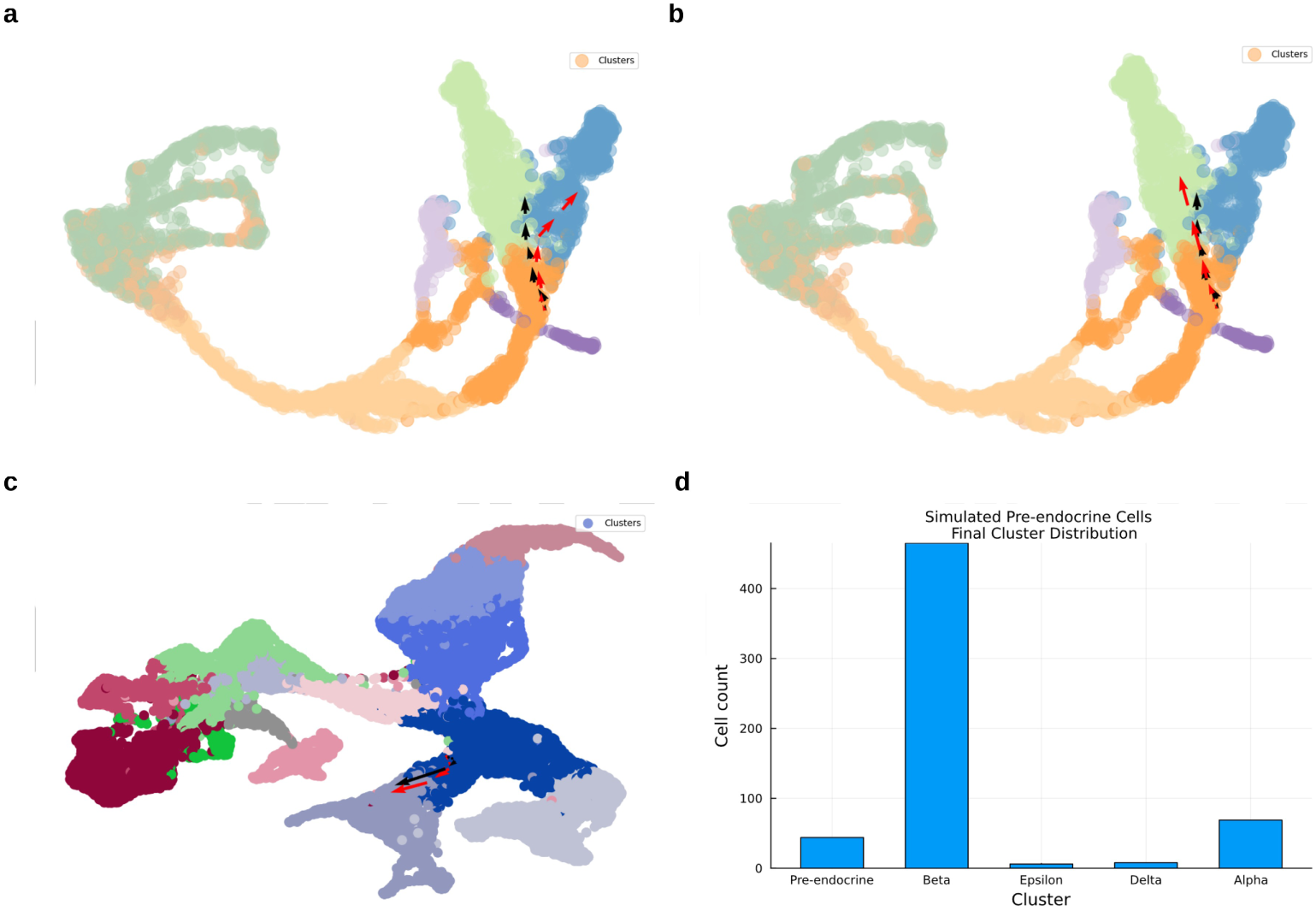
In silico perturbation reveals TF-dependent cell-fate trajectories. **a**. Knockdown of the beta-cell transcription factors Pdx1, Mnx1, and Nr3c1 in pre-endocrine progenitors deflects simulated trajectories away from the beta lineage toward alternative alpha or delta fates. **b**. Knockdown of the alpha-cell drivers Meis2 and Foxp2 produces a comparable lineage disruption. **c**. Knockdown of Zmiz1 in hippocampal ImmGranule1 cells. **d**. Final cluster distribution of simulated pre-endocrine cells without perturbation (baseline); most cells reach the beta-cell cluster.

### JuloVelo demonstrates superior stability and computational efficiency

Finally, we quantitatively benchmarked JuloVelo against scVelo, DeepVelo, and cellDancer across multiple datasets to evaluate its robustness, accuracy, and computational scalability (Fig. 9). Across six independent biological systems, JuloVelo consistently achieved the highest and most stable intrinsic stability scores (quantifying local velocity vector coherence) and extrinsic stability scores (measuring velocity alignment across cell-type transitions) (Fig. 9a–g). In the erythroid gastrulation dataset, JuloVelo was the only method that maintained positive extrinsic stability across all transition boundaries, whereas scVelo and DeepVelo exhibited negative scores due to their incorrect flow directionalities (Fig. 9a). For the Mouse E18 brain dataset, multiome mode further elevated both stability metrics beyond the RNA-only performance, showcasing the quantitative advantage of multi-omic processing (Fig. 9g). In terms of computational efficiency, JuloVelo completed the analysis of the mouse hippocampus dataset (18,213 cells, 2,000 genes) in approximately 10 minutes on an NVIDIA RTX4080 GPU, compared to 28 minutes for cellDancer on an Intel Core i7-13700 CPU, representing a 70% reduction in execution time (Fig. 9h). While the constant-kinetics scVelo and deep-ODE-based DeepVelo were slightly faster (∼4 and ∼8 minutes, respectively), both produced qualitatively incorrect trajectories in complex developmental systems. Thus, JuloVelo provides the optimal trade-off between biophysical modeling accuracy and computational scalability, making it uniquely suited for large-scale single-cell and multi-omic atlas analyses.

**Fig. 9.**
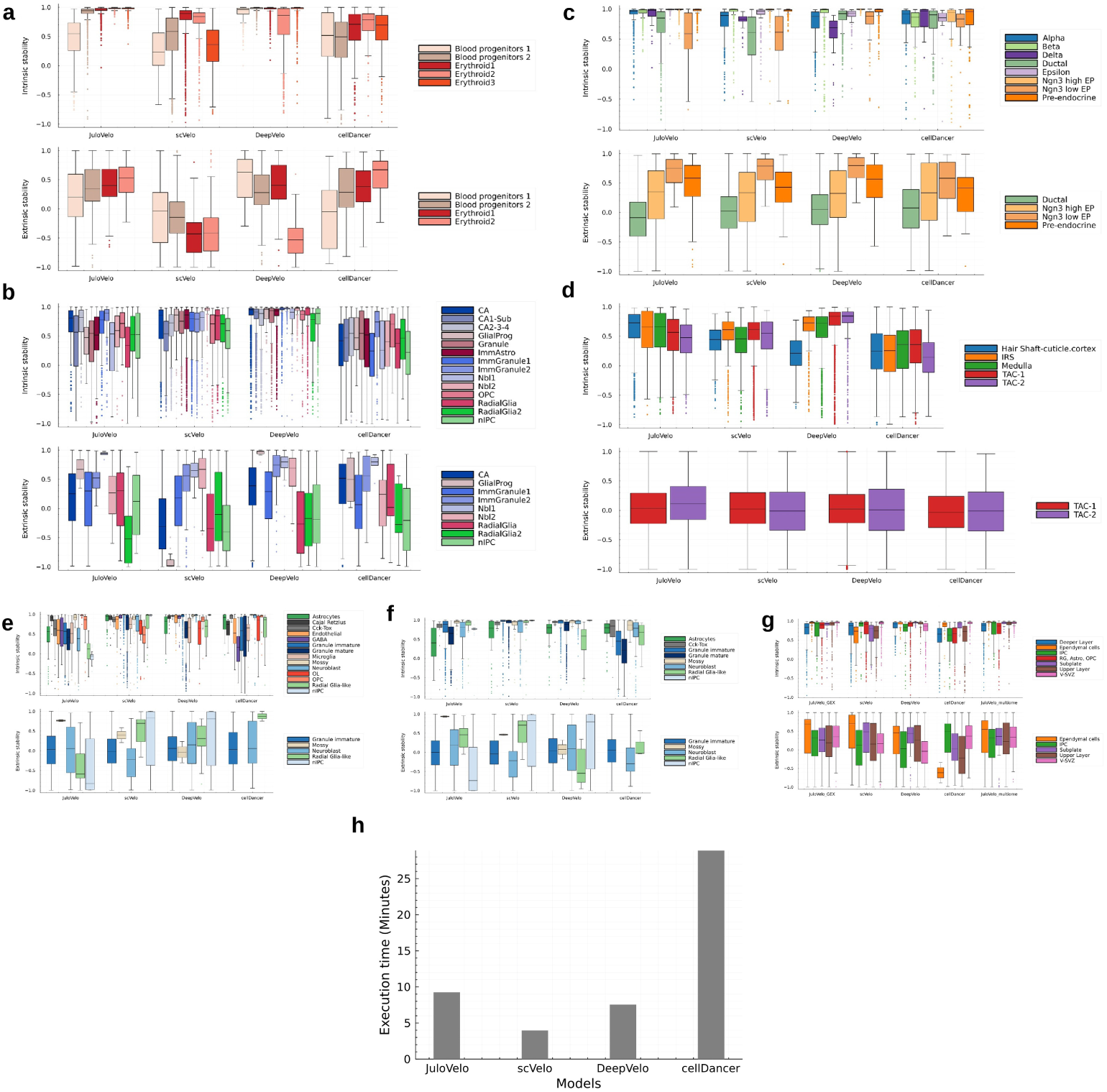
JuloVelo demonstrates superior stability and computational efficiency. Across multiple biological systems, JuloVelo consistently achieves the highest intrinsic (within cell-type) and extrinsic (across boundaries) velocity-stability scores, whereas scVelo and DeepVelo exhibit negative extrinsic scores driven by incorrect flow directionalities. In each panel, intrinsic stability is shown on top and extrinsic stability at the bottom for **a**, mouse erythroid gastrulation; **b**, mouse hippocampus neurogenesis; **c**, mouse pancreatic endocrinogenesis; **d**, mouse skin and hair follicle; **e**, mouse dentate gyrus neurogenesis; **f**, mouse dentate gyrus neurogenesis after removing isolated cell populations; and **g**, mouse embryonic E18 brain, including JuloVelo in RNA-only (GEX) and multiome modes. **h**, Execution time on the 18,213-cell mouse hippocampus dataset: JuloVelo completes the analysis in ∼10 minutes on an NVIDIA RTX4080 GPU, a ∼70% reduction relative to cellDancer (scVelo, DeepVelo, and cellDancer were run on an Intel Core i7-13700 CPU).

## 3 Discussion

In this study, we developed JuloVelo, a deep learning framework that addresses a fundamental gap in RNA velocity inference by introducing gene kinetics trajectory identification as an explicit biological prior for neural network training. By classifying each gene into Circle, Complete induction, or Complete repression patterns based on its unspliced–spliced RNA relationship across cellular transitions, and initializing training with pattern-matched pre-trained networks, JuloVelo provides the cell-specific inference flexibility of cellDancer [12] with substantially improved accuracy. Validated across eight diverse biological datasets, this framework consistently recovered more accurate, continuous differentiation flows and developmental orderings than scVelo [8], DeepVelo [11], and cellDancer [12], particularly in kinetically complex developmental lineages where existing methods fail qualitatively. Furthermore, JuloVelo demonstrates that single-cell multiome datasets can be successfully processed and modeled by using promoter chromatin accessibility to guide gene kinetics predetermination, which resolves kinetics ambiguity and significantly improves velocity stability.

A central advantage of JuloVelo over constant-kinetics methods such as scVelo and UniTVelo [9] is its capacity to model the heterogeneous, time-varying regulatory dynamics that characterize real differentiation processes. Constant kinetic parameters cannot represent transcriptional boosting—the sharp mid-process surge in transcription rate observed for *Hba-x* during erythroid maturation (Fig. 3d,f)—or the simultaneous maintenance of five distinct kinetic states across hippocampal neurogenesis lineages (Fig. 4a,b). Compared to existing cell-specific methods including DeepVelo and cellDancer, JuloVelo’s kinetics pre-determination provides a biologically grounded initialization that reduces convergence to kinetically inconsistent parameter regimes [10]. This guided constraint is most consequential in datasets with highly complex branch topologies or sharp kinetic transitions, where unguided initialization consistently leads to chaotic or reversed velocity vectors.

The cell-specific transcriptional rates inferred by JuloVelo also enable a powerful form of downstream functional gene analysis that is completely unavailable from constant-kinetics models. Conventional approaches to characterizing cell-type-specific function rely on differential expression of genes (DEGs) and downstream pathway enrichment using tools such as Gene Ontology (GO) [17] and KEGG pathways [18], implemented with methods such as DESeq2 [19], edgeR [20], or limma [21]. However, genes in a transcriptional repression state—such as *Rspo3* in erythroid progenitors (Fig. 3c–f) and *Actn4* in pre-endocrine pancreatic cells (Fig. 5e,f)—remain highly detectable by steady-state mature expression long after active transcription has ceased. Treating such genes as active markers contaminates functional enrichment. Differential transcriptional rate analysis using JuloVelo’s cell-specific α estimates (DTRGs) more faithfully represents active, ongoing gene regulatory programs, yielding substantially higher specificity and statistical significance in downstream developmental pathway analyses (Fig. 6).

Despite these advantages, JuloVelo has several limitations that warrant attention. Its local, neighbor-based velocity inference can produce chaotic or undirected flow when applied to isolated or highly discrete cell populations, as observed in mouse dentate gyrus neurogenesis where removing disconnected populations significantly improved vector coherence (Supplementary Figs. S7 and S8). This reflects a general constraint of local velocity models: when the local neighborhood does not include transitional intermediates leading to the expected downstream state, the inferred velocity direction becomes unreliable. A related limitation shared by all splicing-based velocity methods is sensitivity to the quality and completeness of gene annotations. For instance, in the mouse gastrulation erythroid dataset, *Aff3* exhibits anomalously high apparent unspliced counts consistent with an implausible repression state. We attribute this to the recently discovered nested intronic gene *Aff3ir* [22], whose mature transcripts are inadvertently captured as *Aff3* unspliced counts under current annotation schemes, introducing a kinetically inconsistent signal indistinguishable from genuine dynamics.

Looking forward, several extensions of JuloVelo are highly promising. Incorporating global gene status information, such as chromatin accessibility or transcription factor regulatory networks surveyed across all cell types, could stabilize velocity inference for rare or isolated populations that challenge local neighbor-based modeling.

Extending the *in silico* perturbation framework to support combinatorial transcription factor knockdowns and time-resolved trajectory forecasting would greatly broaden its utility for dissecting and engineering complex gene regulatory networks. Additionally, integration with protein-level or spatially resolved transcriptomic measurements could further constrain kinetics inference in tissues where RNA dynamics alone are spatially ambiguous. In conclusion, JuloVelo establishes gene kinetics trajectory identification as a principled, biophysically constrained strategy for cell-specific RNA velocity inference. By bridging deep learning, single-cell kinetics, and multi-omic data modeling, JuloVelo provides a comprehensive computational framework for characterizing and predicting the dynamic regulatory landscapes underlying cell fate decisions.

## 4 Methods

### Overview of JuloVelo

JuloVelo employs deep neural networks (DNNs) to measure cell-specific kinetics and compute RNA velocity, departing from conventional Ordinary Differential Equation (ODE) solver methods. The framework optimizes the network using a customized cosine similarity loss function, assessing the alignment between predicted velocity vectors and the directions towards neighbor cells across genes. The workflow begins by aggregating datasets containing unspliced/spliced RNA counts and optional chromatin accessibility information. Initial processing involves cell and gene filtration, highly variable gene identification, and imputation of RNA counts using scVelo. Subsequently, Leiden clustering and pseudo-time are computed using scanpy. Genes showing low correlation between unspliced and spliced RNA are filtered out. Gene kinetics patterns are pre-determined, guiding the assignment of pre-trained neural networks. Finally, JuloVelo estimates RNA velocity per cell, computes cell velocity in the embedding space, and restores pseudo-time.

The traditional RNA velocity model describes the dynamics of unspliced (u) and spliced (s) RNA for a single gene using ODEs:

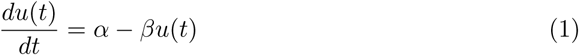

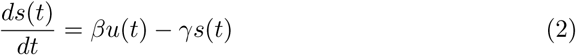

where α, β, and γ represent the constant rates of transcription, splicing, and degradation, respectively. However, recognizing that kinetics can be heterogeneous during differentiation, JuloVelo proposes cell-specific kinetics, modifying the ODEs such that the rates become time-dependent functions:

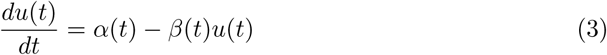

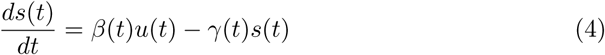

Solving the time-dependent ODEs (3) and (4) analytically or using traditional numerical methods like the Expectation-Maximization algorithm used for constant-rate models is challenging due to the complexity and lack of explicit expressions for α(t), β(t), and γ(t). JuloVelo overcomes this by employing a DNN to approximate these time-dependent functions. For each gene i, a distinct DNN with parameters θ*_i_* takes the unspliced (u*_i_*(t)) and spliced (s*_i_*(t)) RNA amounts as input and outputs the corresponding kinetic parameters:

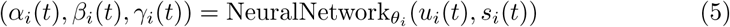

Each gene-specific DNN consists of an input layer (2 nodes), two dense hidden layers (100 nodes each) with Leaky ReLU activation, and an output layer (3 nodes) with Sigmoid activation. The Leaky ReLU activation is defined as:

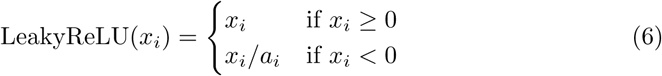

and the Sigmoid activation ensures outputs are within [0, 1]:

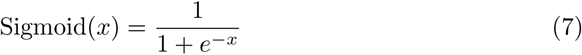

These individual DNNs are effectively merged in parallel into a single 3D network structure for simultaneous training. To mitigate variability from random initialization and improve accuracy, three pre-trained DNNs corresponding to common kinetic patterns (“Circle”, “Complete induction”, “Complete repression”) are utilized based on pre-determined gene kinetics status.

The ODEs (3) and (4) are discretized for the loss calculation, assuming a small time step Δt:

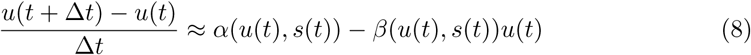

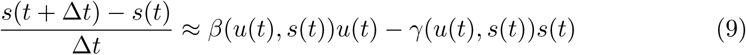

The loss function combines cosine similarity and an L2 penalty. Based on the principle of continuous cell differentiation, a cell’s future state is expected to align with the states of its neighbors. The cosine similarity loss optimizes the alignment between the predicted velocity vector v⃗*_gc_* for gene g in cell c and the state change vector v⃗^nei^ towards its neighbor cells. The loss for cell c and gene g is:

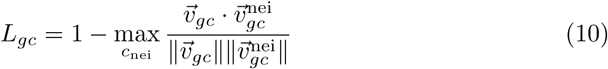

where the velocity vector and neighbor direction vector are:

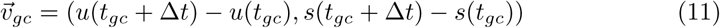

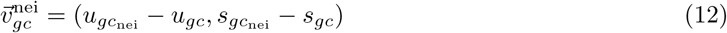

The total cosine loss L_cos_ is the sum over all genes and cells:

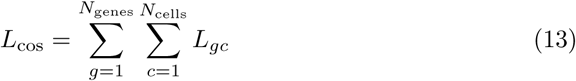

The computation involves finding the k-nearest neighbors (default k=30) for each cell based on its (u, s) state, predicting the velocity vector v⃗*_gc_*, computing the direction vectors to all neighbors v⃗^nei^_gc_, finding the maximum cosine similarity, and summing the (1 *−* max(similarity)) values. To prevent overfitting, L2 regularization is applied to the weights (w*_lcr_*) of the neural network layers:

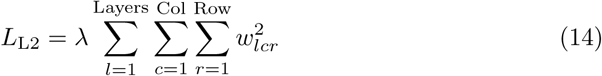

where λ is a hyperparameter. The final loss is a combination of L_cos_ and L_L2_.

For all datasets, training was performed on a default sample of 2400 cells. The Adam optimizer was used with a fixed learning rate of 0.0001. The time shift Δt for velocity calculation was set to 0.5, and the L2 penalty hyperparameter λ was set to 0.004. The network was trained for 100 epochs.

### Dataset Resources

The mouse erythroid gastrulation and mouse gastrulation subset (E7.5) datasets [23] are available from ArrayExpress under accession E-MTAB-6967. The mouse hippocampus neurogenesis, mouse pancreatic endocrinogenesis, and mouse dentate gyrus neurogenesis datasets are available from the NCBI Gene Expression Omnibus (GEO) repository with their accession numbers GSE104323, GSE132188, and GSE95753, respectively. The Human bone marrow dataset is available through the Human Cell Atlas (HCA) data explorer; only its RNA modality was used in this study. In this work, we downloaded the mouse erythroid gastrulation, mouse hippocampus neurogenesis, mouse pancreatic endocrinogenesis, mouse dentate gyrus neurogenesis, human bone marrow, and mouse gastrulation subset to E7.5 data using the scVelo package. The mouse skin and follicle multiome dataset is available on GEO under accession number GSE140203. We downloaded this data from the MultiVelo GitHub page. The Mouse E18 brain multiome dataset is available on the 10x Genomics dataset website. We downloaded this data from the MultiVelo GitHub page.

### Data preprocessing

#### Gene Expression (GEX) Datasets

For GEX datasets, we first assessed the proportion of unspliced and spliced RNA per cell type. Genes with fewer than 20 combined unspliced and spliced reads were filtered out. Read counts were normalized per cell by total counts across all genes, and the top 2000 highly variable genes were selected. Both unspliced and spliced counts were then log-transformed. These steps were performed using scvelo.pp.filter and normalize(adata, min shared counts = 20, n top genes = 2000). The unspliced and spliced proportions of all eight datasets are shown in Supplementary Fig. S9; the multiome datasets (mouse skin and hair follicle, mouse E18 brain, and human bone marrow) contained markedly higher unspliced proportions (47–58%) than the GEX-only datasets (5–25%).

Principal component analysis (PCA) was applied for dimensionality reduction, retaining 30 principal components (PCs). A neighbor graph was computed using 30 nearest neighbors based on Euclidean distance in PCA space. First-order moments for unspliced and spliced matrices were calculated using this neighbor graph information via scvelo.pp.moments(adata, n pcs = 30, n neighbors = 30). If existing dimensionality reduction embeddings (e.g., t-SNE, UMAP) were available in the dataset, they were used directly. Otherwise, UMAP embeddings were generated using scvelo.tl.umap().

#### Multiome Datasets

For multiome datasets (Mouse skin and hair follicle, Mouse E18 brain), RNA counts and chromatin peak counts were extracted from the raw count matrix. Chromatin peak counts underwent basic filtering and imputation using SCALE (SCALE.py -d ATAC.h5ad --min peaks 100 --min cells 3 --n feature -1 --impute).

The RNA counts were pre-processed similarly to the GEX datasets: assessment of unspliced/spliced proportions, gene filtering (minimum 20 shared counts), normalization by total counts per cell, selection of 2000 highly variable genes, and log-transformation of counts using scvelo.pp.filter and normalize(adata, min shared counts = 20, n top genes = 2000).

Chromatin peaks were filtered to retain only those within gene promoter regions. Genes lacking corresponding promoter peaks were removed. The intersection of cells and genes present in both RNA and filtered chromatin peak data was retained. Chromatin peak counts were then log-transformed using scanpy.pp.log1p().

Dimensionality reduction (PCA, 30 PCs) and neighbor graph computation (30 neighbors) were performed on the RNA matrix, followed by moment calculation using scvelo.pp.moments(adata, n pcs = 30, n neighbors = 30). Existing or newly generated UMAP embeddings were used for visualization.

### Clustering and Pseudo-time

Cell clustering was performed using the Leiden algorithm [24], utilizing the nearest neighbor distance matrix and the neighbor graph computed during pre-processing. This step was executed using the scanpy function scanpy.tl.leiden() [25].

To compute pseudo-time, a root cell within a designated root cluster is required [26]. The root cluster was manually determined by integrating cell type annotation with the Leiden cluster information to identify the cluster corresponding to the earliest developmental stage. A representative cell within this root cluster was selected as the root cell using the assignment adata.uns[`’`iroot`’`] = np.flatnonzero(adata.obs[`’`leiden`’`] == root cluster)[0]. Pseudo-time was then calculated using the scanpy function scanpy.tl.dpt() [25].

### Velocity Gene Selection

RNA velocity models infer cell differentiation flow based on modeling gene kinetics. Therefore, unclear or ambiguous gene kinetics patterns can negatively impact the inference accuracy. To mitigate this, we implemented a gene selection step. First, the amounts of unspliced and spliced RNA for each gene were normalized using Min-Max normalization:

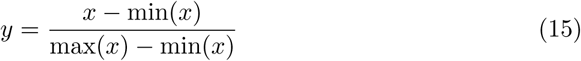

where x represents the original counts and y represents the normalized value.

Next, we calculated the Spearman’s rank correlation coefficient (r*_s_*) between the normalized unspliced and spliced RNA amounts for each gene:

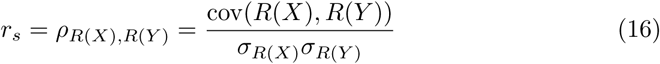

Here, ρ is the standard Pearson correlation coefficient applied to the rank variables R(X) and R(Y ) (ranks of unspliced and spliced RNA amounts, respectively), cov(R(X), R(Y )) is the covariance of the rank variables, and σ*_R_*_(*X*)_ and σ*_R_*_(*Y*)_ are the standard deviations of the rank variables. Genes with a correlation coefficient less than a specified threshold (default set to 0.5) were removed from further analysis.

### Gene Kinetics Trajectory Identification

Single-cell sequencing captures snapshots, meaning not all genes may exhibit their complete kinetic process (induction and repression) within a given sample. We hypothesized that some genes might only be observed in an induction or repression phase. Therefore, we categorized gene kinetics into three types: “Circle” (exhibiting both induction and repression), “Complete induction” (only induction state observed), and “Complete repression” (only repression state observed).

To determine the kinetic category for each gene considered for velocity analysis, we employed different strategies for RNA-seq and Multiome datasets. Recognizing that cells within the same annotated type might have different kinetic statuses, the analysis was performed by partitioning cells based on Leiden clusters, and the overall kinetics for a gene was determined based on the consensus across informative clusters. The root cluster (identified during pseudo-time calculation) was excluded due to potential instability.

For RNA-seq datasets, we first computed the Spearman’s rank correlation between unspliced and spliced RNA amounts within each cluster (excluding the root cluster). Clusters with an absolute correlation below a threshold (default 0.3) were disregarded. For the remaining clusters, we performed linear regression of pseudo-time against unspliced RNA amounts and spliced RNA amounts separately. Clusters were considered informative only if the absolute slopes of both regressions were greater than 0.2 and the ratio of the two slopes was positive. An informative cluster was determined to be in an “Induction” state if both slopes were positive, and in a “Repression” state if both slopes were negative. A gene was categorized as “Complete induction” if all its informative clusters showed an “Induction” state, “Complete repression” if all showed a “Repression” state, and “Circle” if both “Induction” and “Repression” states were observed across its informative clusters.

For Multiome datasets, we incorporated chromatin accessibility information, based on the principle that active transcription often correlates with open chromatin at promoter regions [27]. In addition to the steps performed for RNA-seq data, we computed a linear regression between promoter chromatin accessibility and unspliced RNA amounts for each cluster. A cluster was considered informative only if it passed the RNA-seq criteria AND the absolute slope or intercept of the chromatin accessibility vs. unspliced RNA regression was larger than 0.2. The final gene kinetic categorization (“Complete induction”, “Complete repression”, or “Circle”) followed the same logic as for RNA-seq data, based on the states determined for the informative clusters identified using both RNA and chromatin accessibility criteria.

The overall logic is summarized as follows:

1. **Input**: Root cluster, data type (GEX or Multiome), unspliced/spliced RNA quantities, pseudo-time, and optionally chromatin accessibility.
2. **Initialize**: Empty list cluster kinetics.
3. **For each cluster:**

- Calculate Spearman correlation (us) between unspliced and spliced RNA.
- Calculate linear regression slopes: tu (pseudo-time vs. unspliced) and ts (pseudo-time vs. spliced).
- For Multiome: calculate cu (chromatin accessibility vs. unspliced).
- Skip cluster if *|*us*|* < 0.3.
- Skip cluster if *|*tu*|* < 0.2, *|*ts*|* < 0.2, or tu/ts < 0.
- For Multiome: skip cluster if cu.slope < 0.2 and cu.intercept < 0.2.
- Skip if cluster is the root cluster.
- If tu > 0: add “Induction” to cluster kinetics.
- If tu < 0: add “Repression” to cluster kinetics.
4. Determine gene kinetics:

- If both “Induction” and “Repression” present: “Circle”.
- If only “Induction” present: “Complete induction”.
- If only “Repression” present: “Complete repression”.
- Otherwise: “fail”.

### Velocity Projection and Visualization

To project the high-dimensional RNA velocity vectors onto lower-dimensional embeddings (like t-SNE or UMAP) for visualization, we first applied a variance-stabilizing transformation (VST) to the velocity v*_ic_* of each gene i in cell c:

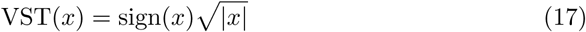

The VST-transformed velocity vector for cell c across all g analyzed genes is then v⃗*_c_* = VST(v_1*c*_, v_2*c*_, . . . , v*_gc_*).

Following the approach of La Manno et al. [7], as implemented in scVelo [8], we project velocities by estimating cell-to-cell transition probabilities. First, the change in gene expression between cell c and a neighbor cell c_nei_ is calculated using VST:

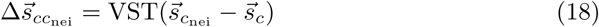

The correlation between the cell’s velocity vector v⃗*_c_* and the expression change vector 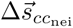 indicates the likelihood of a transition from c to c_nei_. This correlation is computed as the cosine similarity:

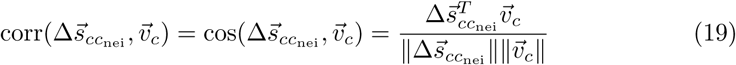

The transition probability P*_cc_*_nei_ is computed using an exponential kernel based on this correlation:

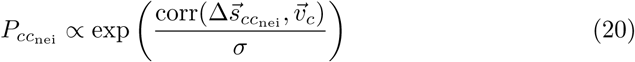

where σ is a kernel width parameter (default 0.05). These probabilities are calculated for the k-nearest neighbors (default k = 200) of cell c in the embedding space. The probabilities are then normalized to sum to 1 across the neighbors:

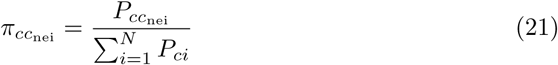

where N is the number of neighbors considered.

Finally, the projected velocity vector v⃗_embedding*,c*_ for cell c in the low-dimensional embedding space (with coordinates x⃗*_c_*) is computed as an expected displacement based on the transition probabilities:

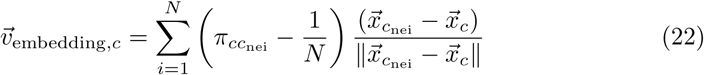

The term 1/N corrects for potential non-uniform density of points in the embedding.

For visualization, the computed velocity information is exported into an AnnData structure using JuloVelo.to anndata() and JuloVelo.write h5ad(). The velocity stream plot on the embedding space is then generated using the scVelo function scvelo.pl.velocity embedding stream().

### Transcription Factor Identification and Perturbation Simulation

To investigate how transcription factor (TF) activity influences cell fate decisions, we developed an *in silico* perturbation framework that combines regulatory network inference with velocity-based simulation.

#### Regulatory Network Inference with pySCENIC

Transcription factors and their target genes (regulons) were identified using pySCENIC (py Single-Cell rEgulatory Network Inference and Clustering). This pipeline consists of three main steps: (1) GRN inference using GENIE3 to identify potential TF-target relationships, (2) regulon prediction via RcisTarget to identify direct targets based on cis-regulatory motifs, and (3) cellular regulon activity scoring using AUCell to quantify regulon activity in individual cells. For each dataset, we selected cell-type-specific TFs based on high regulon activity scores (AUCell) and specificity to particular cell clusters.

#### In Silico Perturbation Design

The perturbation simulation framework leverages the cell-specific kinetics parameters (α, β, γ) inferred by JuloVelo for each gene in each cell. For a selected transcription factor and its regulon (target genes), we implement two perturbation modes:

### Knockdown simulation

For all genes in the regulon, the transcription rate α is set to zero, effectively silencing the transcriptional activity of the TF’s downstream targets. This mimics the effect of TF knockdown or inhibition.

### Overexpression simulation

For all genes in the regulon, the initial unspliced (u_0_) and spliced (s_0_) RNA levels are set to a high quantile (e.g., 90th percentile) of their observed distributions. The transcription rate α may also be scaled up to reflect increased transcriptional activity.

#### Euler Method for State Projection

Following perturbation initialization, we project the cell state forward in time using the Euler method to solve the time-dependent ODEs (3) and (4). Given the current state (u(t), s(t)) and the kinetic parameters (α(t), β(t), γ(t)) predicted by the neural network, the state at the next time step t + Δt is computed as:

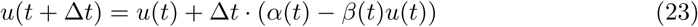

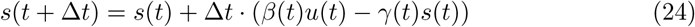

where Δt is a small time step (default 0.5). This process is iterated for multiple steps to simulate the trajectory of the perturbed cell state. The final cluster assignment is determined by projecting the simulated state onto the UMAP embedding and identifying the nearest neighbor cluster.

## Supplementary information

Supplementary Figs. S1–S9 are provided at the end of this manuscript.

## Declarations

### Funding

This work was supported by the National Science and Technology Council, Taiwan (NSTC 113-2320-B-002-025-MY3, NSTC 113-2221-E-002-149-MY3, NSTC 115-2218-E-002-017, NSTC 115-2321-B-002-027, NSTC 115-2221-E-A49-115-MY3), Research Proposal for NTU Core Consortiums (NTU-CC-115L892702 and NTU-115L8509), and the Center for Advanced Computing and Imaging in Biomedicine (NTU-115L900701) from the Featured Areas Research Center Program within the framework of the Higher Education Sprout Project by the Ministry of Education (MOE) in Taiwan. The funders had no role in study design, data collection and analysis, decision to publish, or preparation of the manuscript.

### Competing interests

The authors declare no competing interests.

### Ethics approval and consent to participate

Not applicable. This study used only publicly available, de-identified datasets.

### Data availability

All datasets analysed in this study are publicly available; their accession numbers and sources are listed in the Methods (Dataset Resources).

### Code availability

JuloVelo is implemented in Julia and is freely available under the MIT license at https://github.com/jhlabtw/JuloVelo.jl.

### Author contributions

K.-C.T.: Methodology, Software, Visualization, Writing – original draft. Y.-H.T.: Conceptualization, Methodology, Software, Writing – review & editing. K.-Y.H.: Methodology, Software, Visualization, Validation. H.-F.J.: Supervision, Funding acquisition, Writing – review & editing. H.-C.H.: Conceptualization, Methodology, Supervision, Funding acquisition, Writing – review & editing. All authors read and approved the final manuscript.

---

**Fig. S1.**
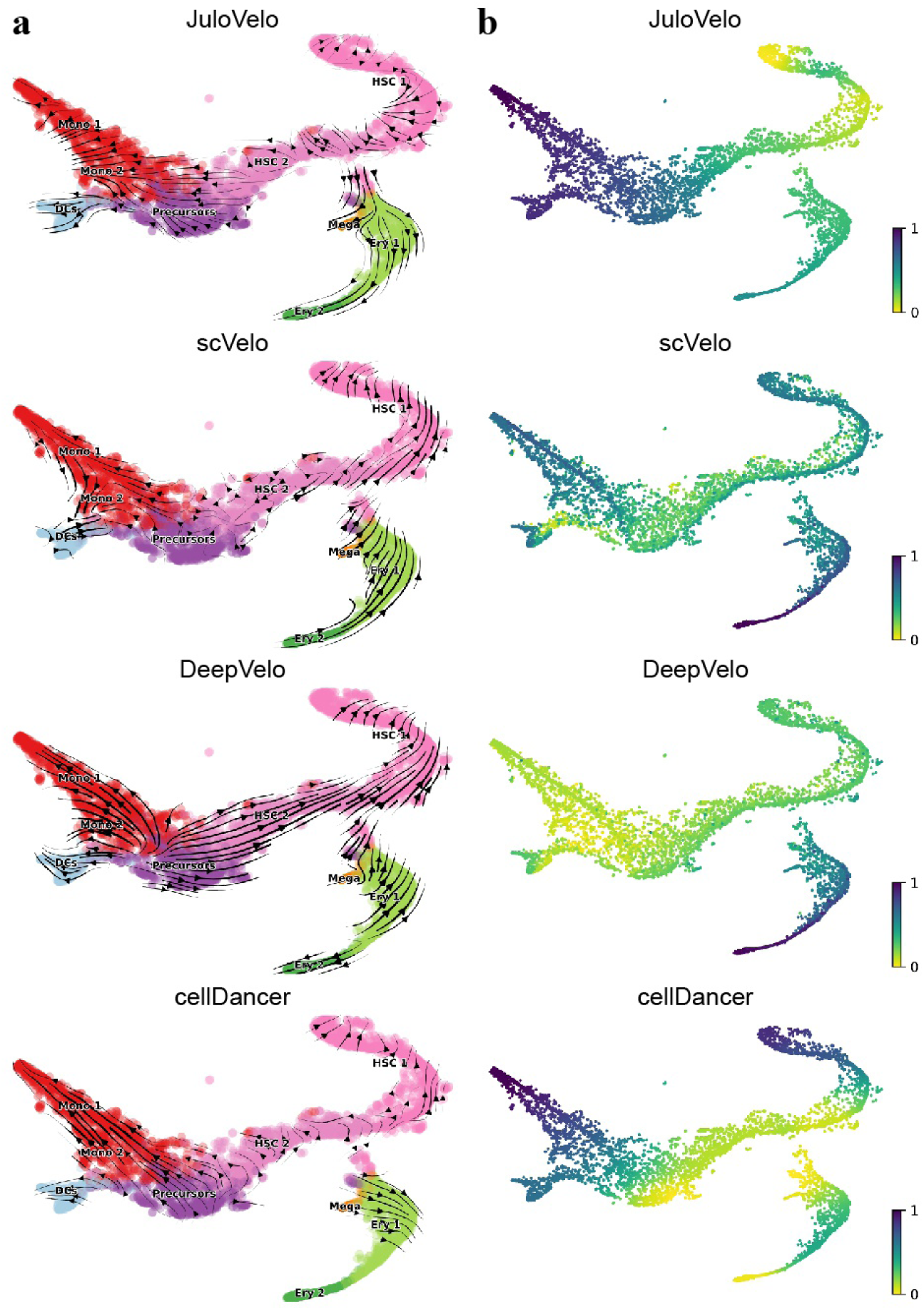
Velocity inference in human bone marrow hematopoiesis. **a**, Velocity streams of JuloVelo, scVelo, DeepVelo, and cellDancer projected on the UMAP embedding. **b**, Velocity pseudotime inferred by each method. All methods show inconsistent flows within HSC_1 and HSC_2, and JuloVelo additionally shows unexpected flows in early HSC_1 and in immune cell types.

**Fig. S2.**
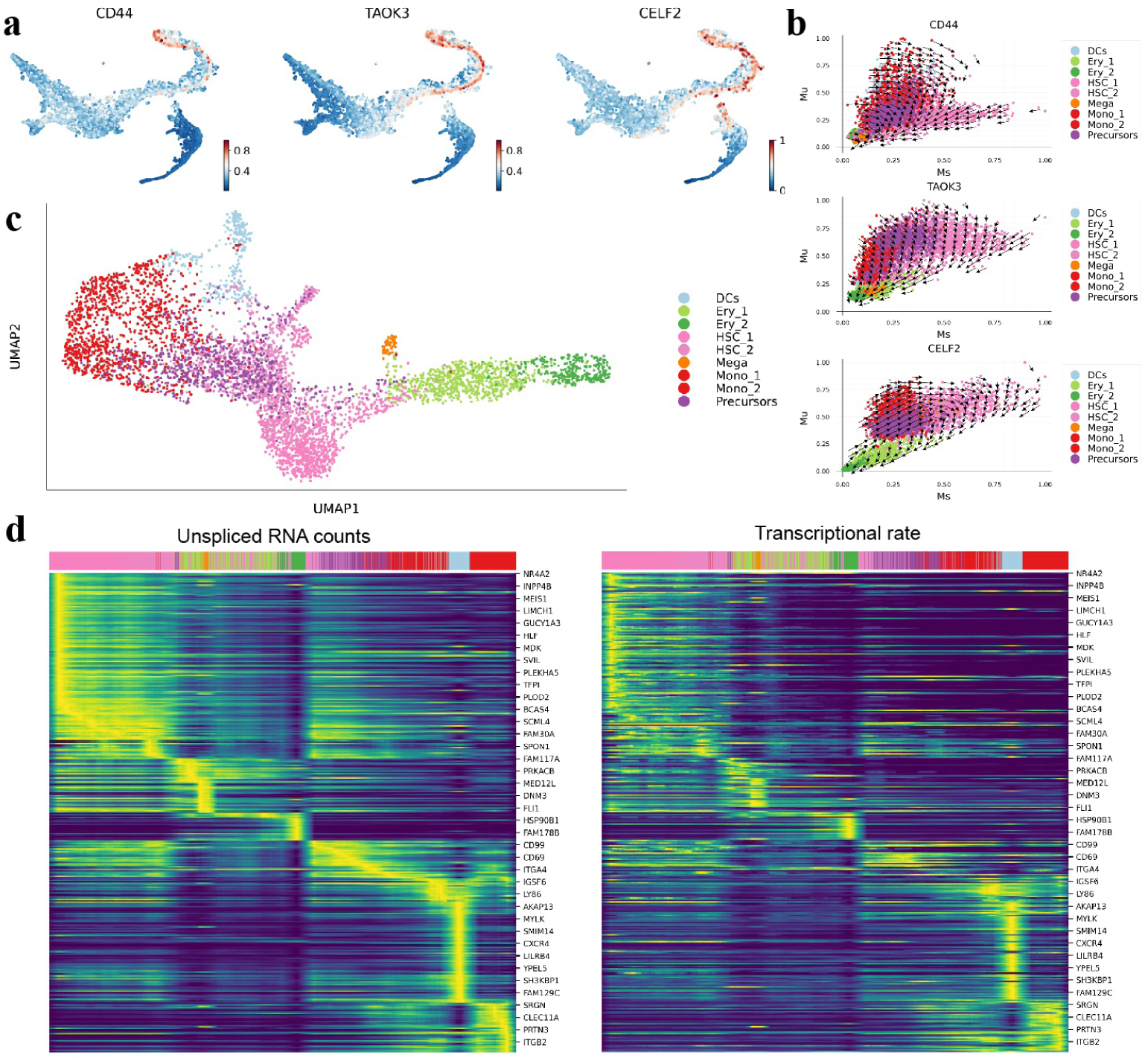
Gene kinetics inferred by JuloVelo in human bone marrow hematopoiesis. **a**, Expression of the HSC-expressed genes *CD44*, *TAOK3*, and *CELF2* on the UMAP embedding. **b**, Unspliced–spliced phase portraits of these genes with JuloVelo velocity vectors, showing opposite velocity directions in HSCs and immune cells. **c**, UMAP embedding constructed from the cell-specific kinetics parameters estimated by JuloVelo. **d**, Heatmaps of unspliced RNA counts (left) and JuloVelo transcriptional rates (right) for velocity genes, with cells ordered by pseudotime.

**Fig. S3.**
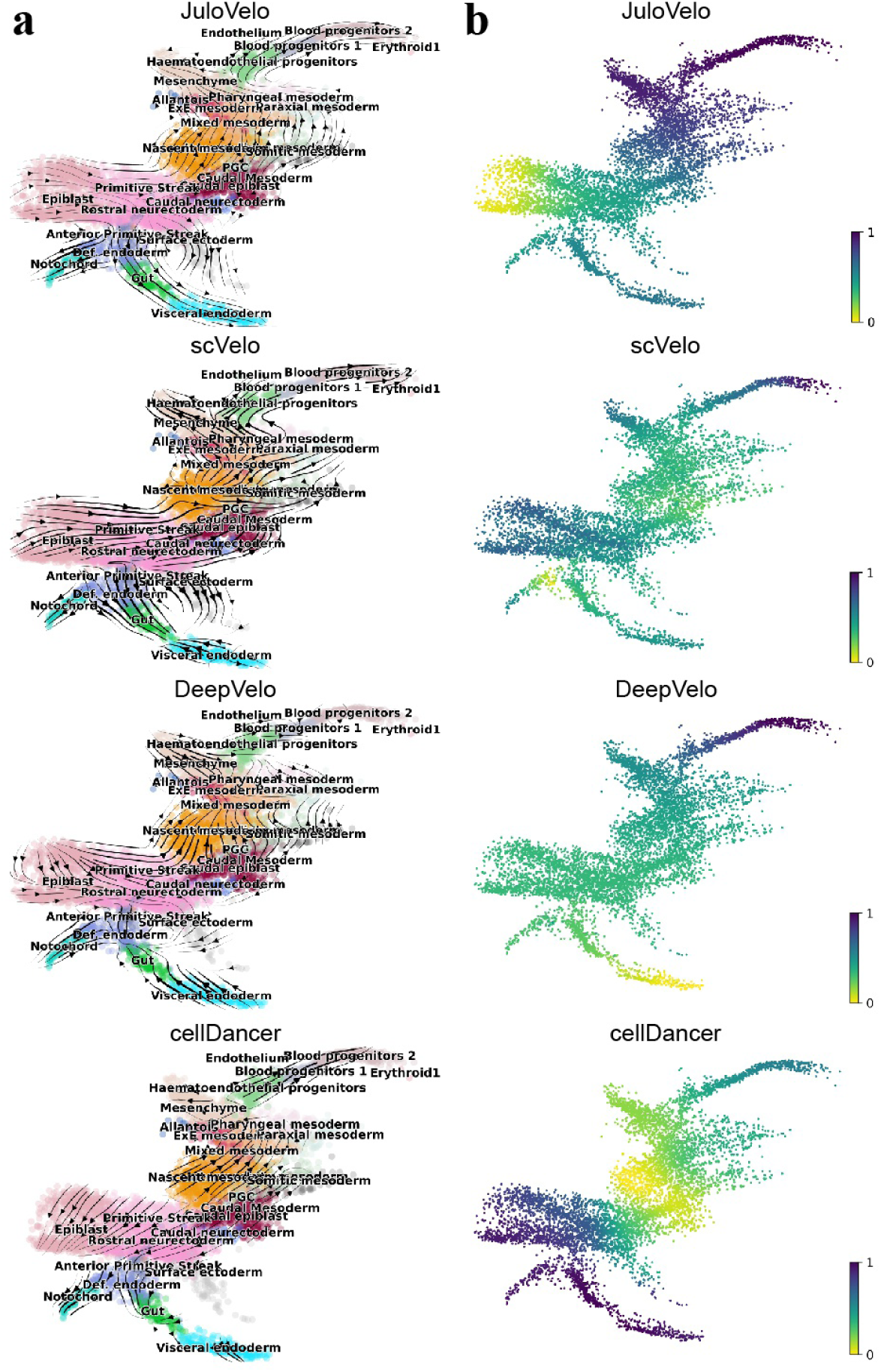
Velocity inference in the mouse gastrulation subset to E7.5. **a**, Velocity streams of JuloVelo, scVelo, DeepVelo, and cellDancer projected on the UMAP embedding. **b**, Velocity pseudotime inferred by each method. Only JuloVelo recovers a reasonable developmental ordering.

**Fig. S4.**
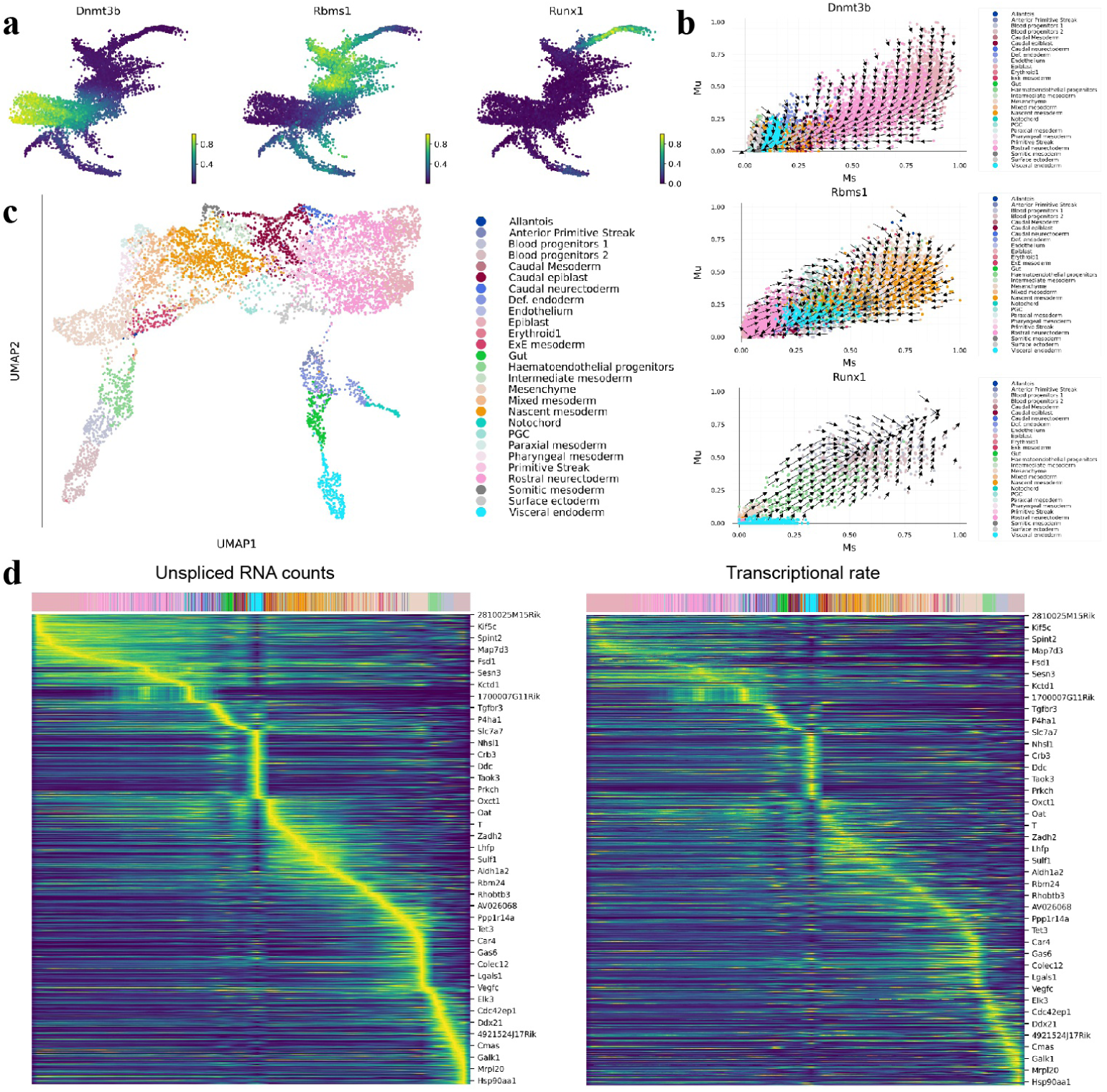
Gene kinetics inferred by JuloVelo in the mouse gastrulation subset to E7.5. **a**, Expression of *Dnmt3b*, *Rbms1*, and *Runx1* on the UMAP embedding. **b**, Unspliced–spliced phase portraits of these genes with JuloVelo velocity vectors; *Runx1* shows induction-only kinetics along erythroid differentiation. **c**, UMAP embedding constructed from the cell-specific kinetics parameters estimated by JuloVelo, which reconstructs the full differentiation process. **d**, Heatmaps of unspliced RNA counts (left) and JuloVelo transcriptional rates (right) for velocity genes, with cells ordered by pseudotime.

**Fig. S5.**
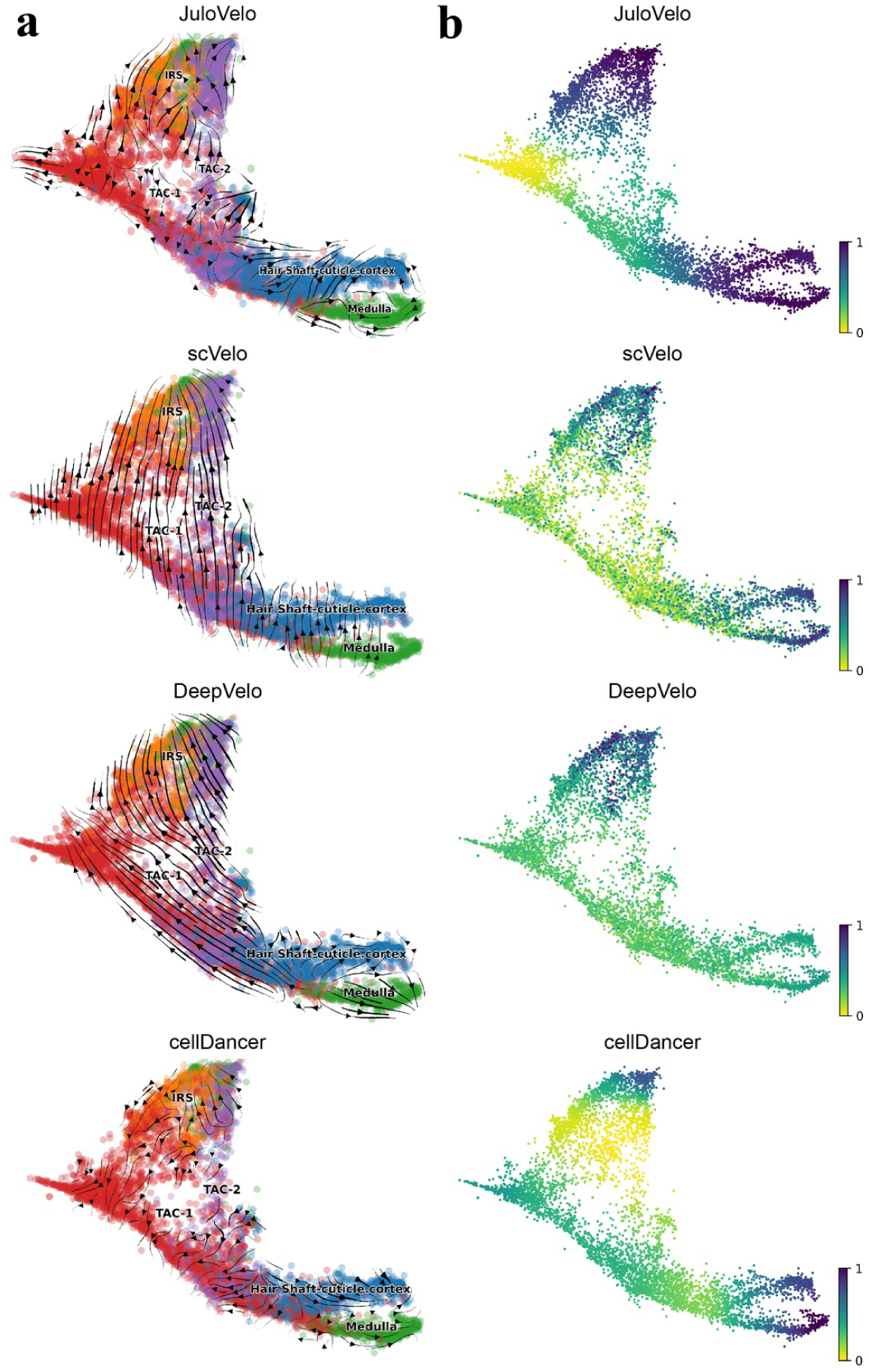
Velocity inference in mouse skin and hair follicle. **a**, Velocity streams of JuloVelo, scVelo, DeepVelo, and cellDancer projected on the UMAP embedding. **b**, Velocity pseudotime inferred by each method. Only JuloVelo recovers a reasonable flow and pseudotime from TAC-1 towards the IRS, hair shaft cuticle and cortex, and medulla, with slightly irregular flow within TAC-1 and TAC-2.

**Fig. S6.**
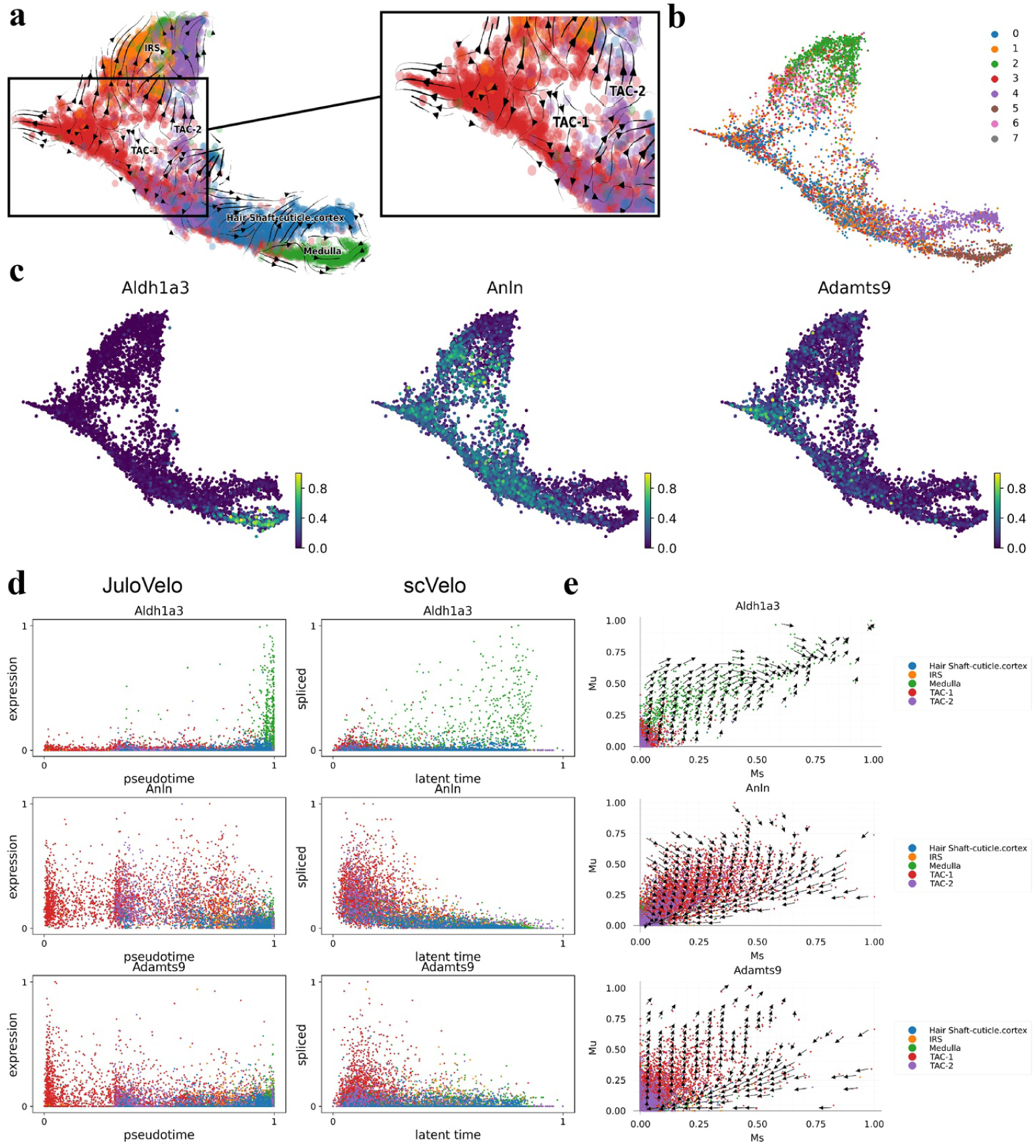
Gene kinetics inferred by JuloVelo in mouse skin and hair follicle. **a**, JuloVelo velocity streams, with the TAC-1 and TAC-2 region enlarged. **b**, Leiden clusters; TAC-1 and TAC-2 each contain several evenly represented clusters. **c**, Expression of *Aldh1a3*, *Anln*, and *Adamts9* on the UMAP embedding. **d**, Expression of these genes along JuloVelo pseudotime (left) and spliced counts along scVelo latent time (right). **e**, Unspliced–spliced phase portraits with JuloVelo velocity vectors.

**Fig. S7.**
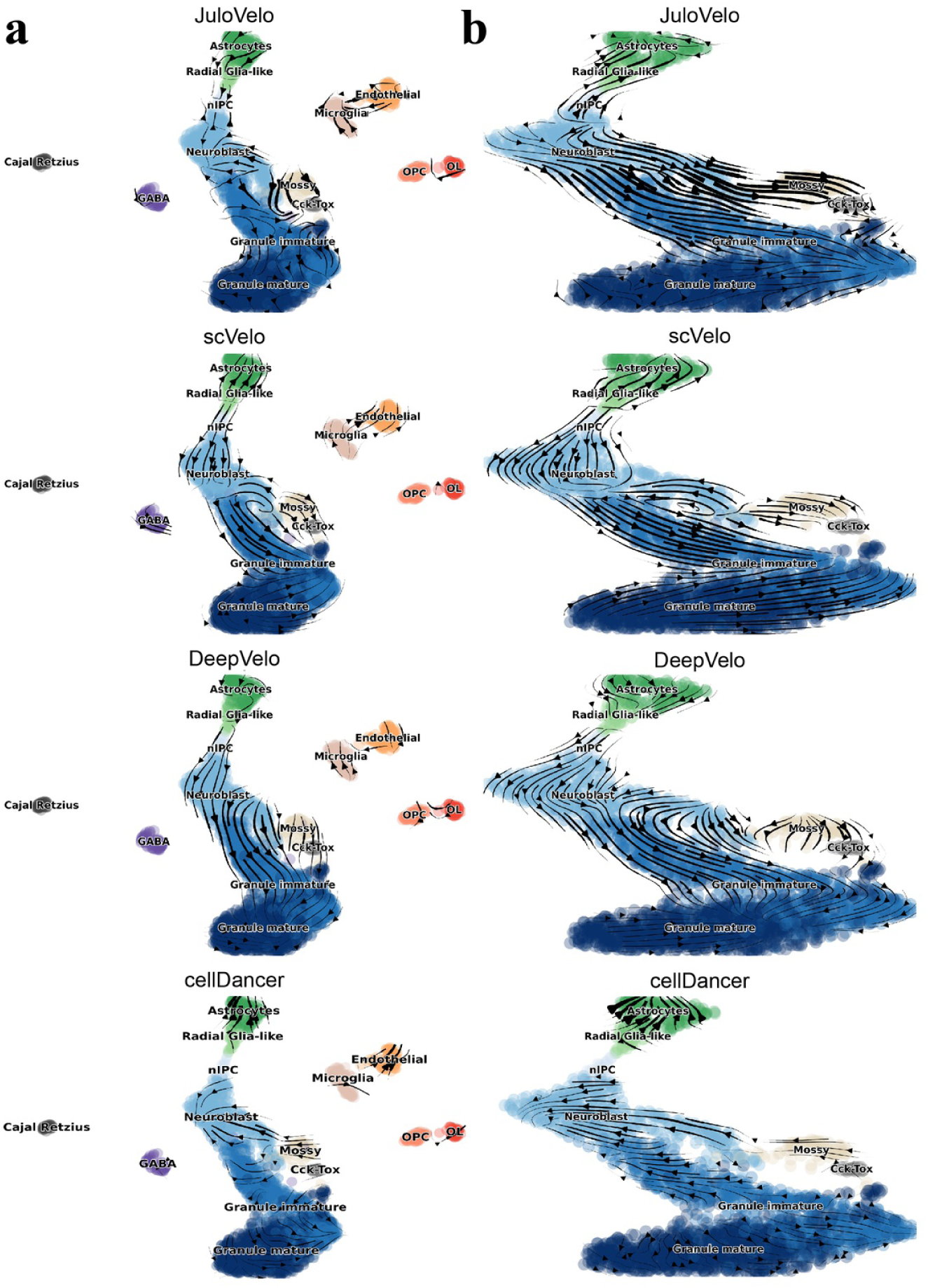
Velocity inference in mouse dentate gyrus neurogenesis. Velocity streams of JuloVelo, scVelo, DeepVelo, and cellDancer on (**a**) the full dataset and (**b**) the main differentiation branch after removing isolated cell populations. JuloVelo and cellDancer show chaotic flows on the full dataset and recover more stable flows after removal; none of the methods correctly infers flow terminating in mature granule cells.

**Fig. S8.**
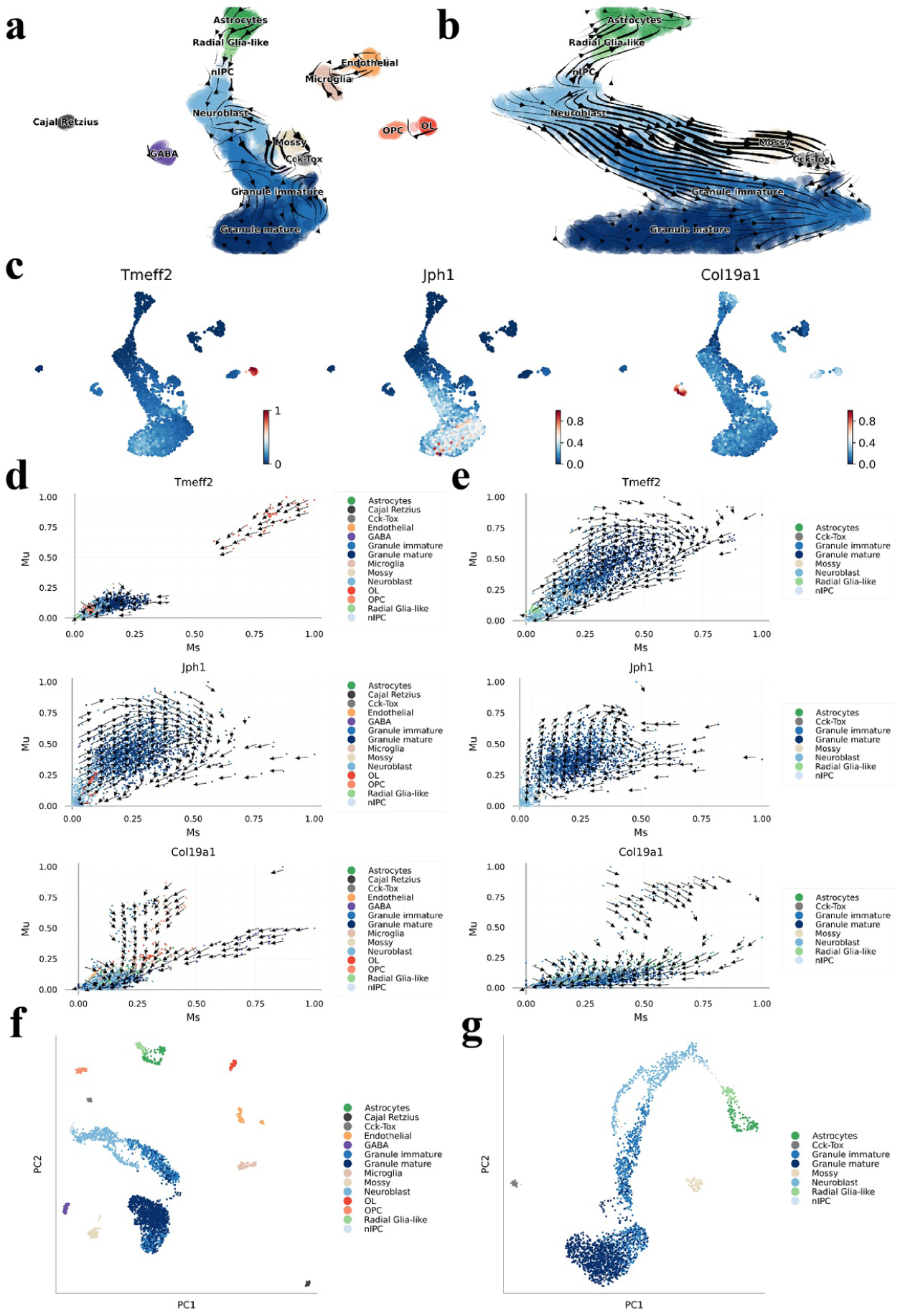
Isolated cell populations affect local velocity inference in mouse dentate gyrus neurogenesis. JuloVelo velocity streams on (**a**) the full dataset and (**b**) the main differentiation branch. **c**, Expression of *Tmeff2*, *Jph1*, and *Col19a1* on the UMAP embedding. Unspliced–spliced phase portraits with JuloVelo velocity vectors for (**d**) the full dataset, where isolated populations such as OL and GABA distort the distribution of the main branch, and (**e**) the main branch only. Embedding of cell-specific kinetics for (**f** ) the full dataset and (**g**) the main branch.

**Fig. S9.**
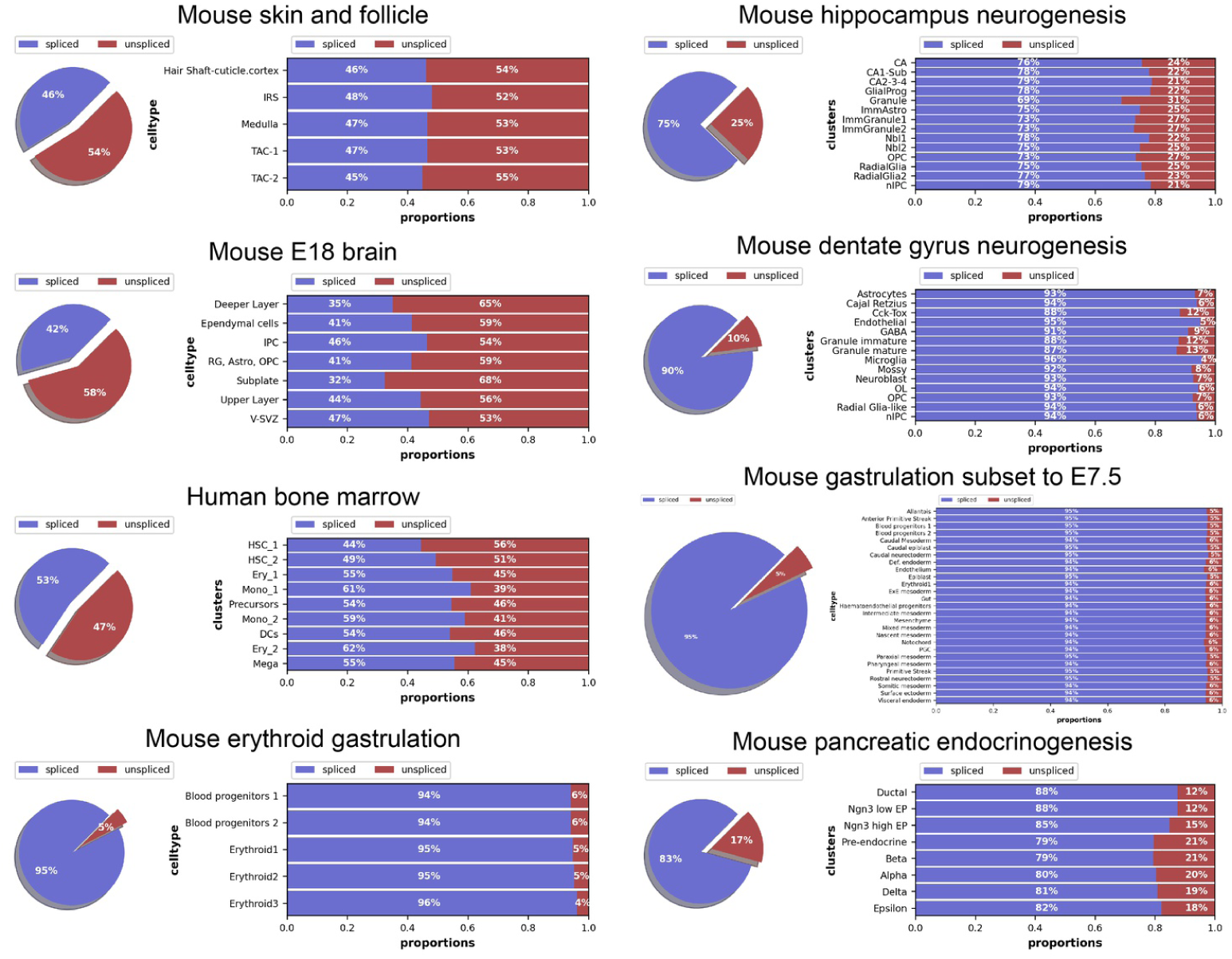
Proportions of spliced and unspliced reads in the eight datasets. For each dataset, the pie chart shows the overall proportions and the bar chart shows the proportions per cell type. The multiome datasets (mouse skin and hair follicle, mouse E18 brain, and human bone marrow) contain markedly higher unspliced proportions than the GEX-only datasets.

## Notes

https://github.com/jhlabtw/JuloVelo.jl

### Competing Interest Statement

The authors have declared no competing interest.

